# Allele-specific alternative splicing in human tissues

**DOI:** 10.1101/2020.05.04.077255

**Authors:** Kofi Amoah, Yun-Hua Esther Hsiao, Jae Hoon Bahn, Yiwei Sun, Christina Burghard, Boon Xin Tan, Ei-Wen Yang, Xinshu Xiao

## Abstract

Alternative splicing is an RNA processing mechanism that affects most genes in human, contributing to disease mechanisms and phenotypic diversity. The regulation of splicing involves an intricate network of *cis*-regulatory elements and *trans*-acting factors. Due to their high sequence specificity, *cis*-regulation of splicing can be altered by genetic variants, significantly affecting splicing outcomes. Recently, multiple methods have been applied to understanding the regulatory effects of genetic variants on splicing. However, it is still challenging to go beyond apparent association to pinpoint functional variants. To fill in this gap, we utilized large-scale datasets of the Genotype-Tissue Expression (GTEx) project to study genetically-modulated alternative splicing (GMAS) via identification of allele-specific splicing events. We demonstrate that GMAS events are shared across tissues and individuals more often than expected by chance, consistent with their genetically driven nature. Moreover, although the allelic bias of GMAS exons varies across samples, the degree of variation is similar across tissues vs. individuals. Thus, genetic background drives the GMAS pattern to a similar degree as tissue-specific splicing mechanisms. Leveraging the genetically driven nature of GMAS, we developed a new method to predict functional splicing-altering variants, built upon a genotype-phenotype concordance model across samples. Complemented by experimental validations, this method predicted >1000 functional variants, many of which may alter RNA-protein interactions. Lastly, 72% of GMAS-associated SNPs were in linkage disequilibrium with GWAS-reported SNPs, and such association was enriched in tissues of relevance for specific traits/diseases. Our study enables a comprehensive view of genetically driven splicing variations in human tissues.

## Introduction

High-throughput sequencing technologies are enabling identification of an extraordinary number of genetic variants in the human genome^1^. These data provide a foundation to elucidate the genetic underpinnings of human diseases or phenotypic traits. Many genome-wide studies have been conducted to uncover associations between the genetic variants and complex traits^2^. However, moving from associations to revealing the underlying mechanisms remains a significant challenge. Genetic variants could affect many aspects of gene expression or function, which is a major determinant of phenotypic diversity^3^. Until recently, research efforts have been focused on variants that may impose epigenetic or transcriptional regulation. However, it is increasingly recognized that genetic variants also play critical roles in modulating post-transcriptional mechanisms, such as alternative splicing^4,5^.

Alternative splicing is an essential mechanism in eukaryotic gene expression, contributing to many aspects of phenotypic complexity and disease mechanisms^6^. Splicing is regulated by an intricate network of *trans*-factors and *cis*-regulatory elements^6^. Thus, it is not surprising that genetic variants may alter different aspects of splicing regulation, such as the *cis*-regulatory motifs, *trans*-factor expression or function, and the interactions between these players^4,5^. Indeed, quantitative trait loci (QTL) mapping in lymphoblastoid cell lines suggested that splicing QTLs and expression QTLs are comparable in their effects on complex traits^7,8^.

Both computational and experimental methods have been developed to reveal splicing-disrupting genetic variants^9,10,11^. Computationally, applications of machine learning methods have yielded promising results^12^. Recently, performance improvements were achieved using deep learning to predict splice site usage directly from nucleotide sequence^13,14,15,16^. However, these methods still present challenges in interpretability and it is difficult to determine whether the features being used are biologically relevant. Experimentally, massively parallel reporter assays have enabled large-scale screens of functional variants in splicing^17,18,19,20,21^. However, due to the limited insert size cloned into the reporters, the splicing outcome may not always recapitulate endogenous splicing patterns. Additionally, these reporter assays can only be performed in one cell type at a time. In general, it remains a great challenge, both computationally and experimentally, to identify causal genetic variants specific to each tissue type.

In this study, we carried out global analyses of allele-specific alternative splicing using RNA-seq data from a large panel of human tissues and individuals generated by the GTEx project^22^. Compared to machine learning methods, allele-specific analysis is a data-driven approach that requires little *prior* knowledge about splicing regulatory mechanisms. The advantage of this approach includes its applicability to a single RNA-seq dataset. In addition, it compares the alternative alleles of a heterozygous SNP in the same cellular environment in the same subject. Thus, the method controls for tissue conditions, *trans*-acting factors, global epigenetic effects, and other environmental influences.

Our lab previously developed allele-specific analysis methods to identify allele-specific splicing events, also called genetically modulated alternative splicing (GMAS)^23,24^. Here, in addition to applying these methods to the GTEx data, we developed a new method to infer functional SNPs underlying the GMAS events. Using these methods, we observed that GMAS patterns were significantly shared between tissues and individuals, consistent with the expectation that genetic variants are the driving factors for GMAS. Nonetheless, some GMAS events showed high variability across individuals or tissues, enriched in genes related to immune response or muscle function, respectively. Importantly, the degree of variability in allele-specific splicing of GMAS exons was similar across tissues and individuals. The large-scale dataset also allowed us to examine the functional relevance of GMAS events. About 72% of GMAS-associated SNPs were in linkage disequilibrium (LD) with GWAS-reported SNPs, a significantly higher percentage than expected by chance. Moreover, for a number of GWAS traits, the related GMAS events were enriched in tissues expected to be closely relevant to the traits.

## Results

### Overview of genetic modulation of alternative splicing in GTEx data

We first applied our previously published method^23^ to identify GMAS events. Briefly, this method examines allelic biases in reads covering heterozygous SNPs in each gene. By comparing the allele-specific expression patterns of all heterozygous SNPs in a gene and their associations with alternative splicing, the method identifies SNPs that are associated with allelespecific splicing patterns (Methods). Although this method does not pinpoint the functional (or causal) SNPs regulating splicing, it captures exons (namely GMAS exons) that are under such genetic regulation. Therefore, the SNPs with allelic bias located in the GMAS exons are named tag SNPs. Using this method, we analyzed a total of 7,822 GTEx RNA-seq datasets, across 47 tissues and 515 donors, following a few quality control filters (Methods).

Across all tissues, a total of 12,331 exons were identified as GMAS exons, associated with 18,894 heterozygous tag SNPs (Methods), where one GMAS exon may be associated with multiple tag SNPs. We focused on GMAS events that are common to multiple samples by requiring each GMAS exon be present in ≥3 samples (across all tissues and individuals). A total of 4,941 GMAS exons (7,404 tag SNPs) were retained (Supplemental Table 1). For each tissue, an average of 10% of all testable exons (defined based on read coverage requirements, see Methods) were identified as GMAS exons (Fig. 1A). This percentage is highest in whole blood (17.8%), which may reflect existence of a high level of genetic modulation of splicing, consistent with the sQTL results in the GTEx study^22^. Pancreas, in which splicing regulation is not well understood^26^, had the smallest fraction of exons demonstrating GMAS patterns (8.4%). This lower percentage could be partly explained by the low read depth relative to other tissues (Supplemental Fig. S1).

**Figure 1:**
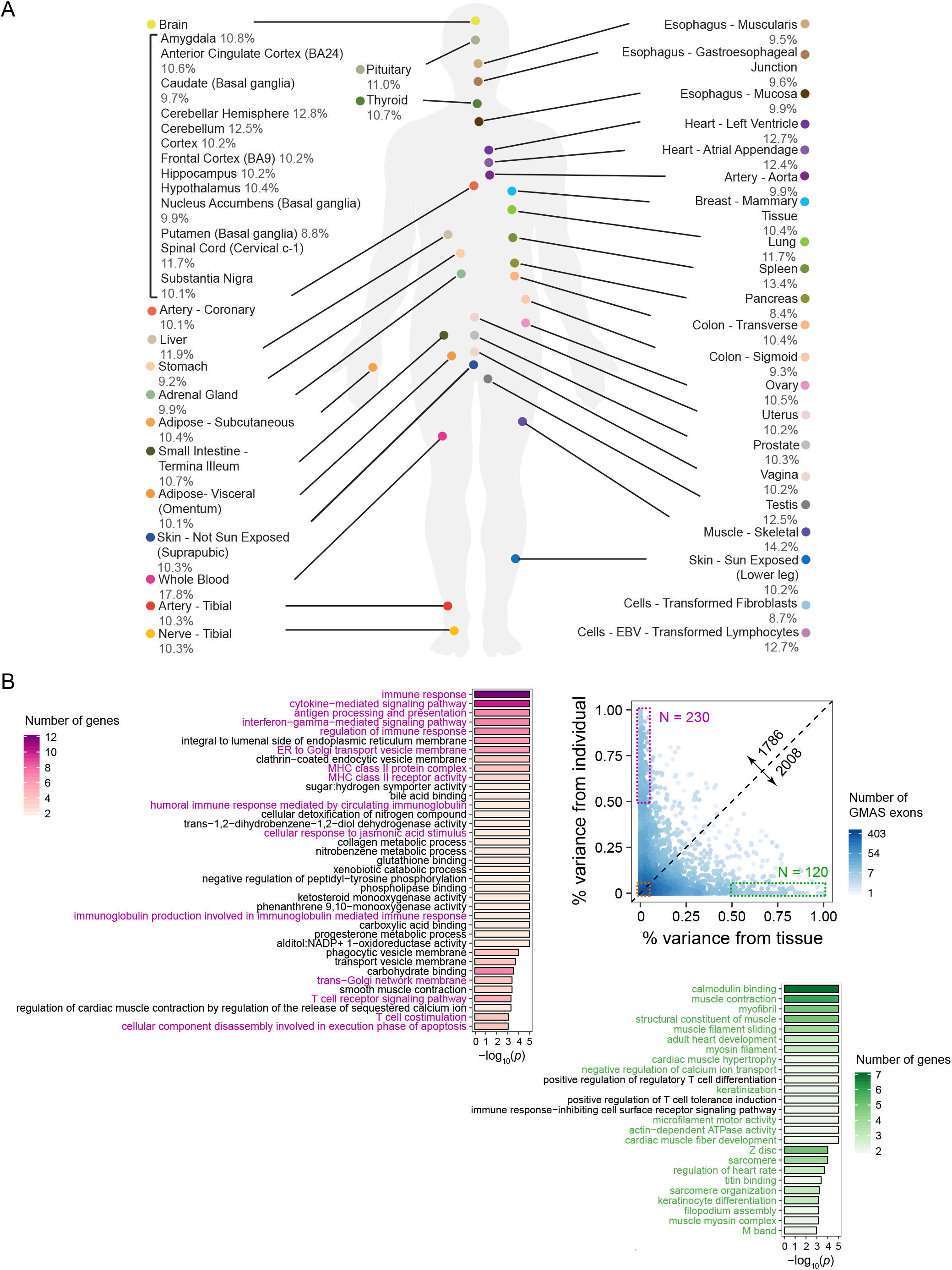
The landscape of GMAS exons in human tissues. (A) % of GMAS exons among all testable exons in each tissue (averaged across individuals). (B) The variability of GMAS patterns across tissues and individuals (Methods). Each dot represents an exon, and the colors represent the number of overlapping dots. This analysis only included GMAS exons that exist in ≥2 individuals per tissue and ≥2 tissues per individual. The numbers along the diagonal line show the number of GMAS exons above and below the line, respectively. GO terms enriched among genes in the high variability groups (boxed) are shown. Color intensity represents the number of genes associated with each significant GO term. The *p* values were estimated based on 10,000 randomizations of control genes matching gene length and GC content of the test genes^24^. The significant cutoff of the *P* value was set to be 1/(number of total GO terms considered).

In each tissue, the most prevalent type of alternative splicing for GMAS events is skipped exons (SEs), accounting for about 80% of all events, followed by retained introns (RIs, ~10%) (Supplemental Fig. S2A). The distribution of percent spliced-in (PSI) values of GMAS exons in each tissue generally showed bimodal patterns, except for the RI events that had relatively low PSI (Supplemental Fig. S2B), consistent with previous findings for alternatively spliced exons in general^27,28^.

### GMAS patterns vary across tissues and individuals to a similar degree

Given the datasets from many individuals and a large panel of tissues, we first examined the global variability in GMAS patterns depending on these two variables. To segregate the impact of tissues and individuals on GMAS, we used a linear mixed model that includes these two variables and a number of confounding factors (age, ethnicity and gender) (Methods). We observed equivalent levels of dependence of GMAS on tissues and individuals (Fig. 1B). This result is in stark contrast to previous findings that both gene expression and splicing in general predominantly vary depending on tissue types instead of individuals^29^. Nevertheless, our result is not surprising because GMAS, by definition, consist of splicing events modulated by genetic variants that can be individual-specific. In turn, this result confirms the validity of the reported GMAS events. Importantly, our observation highlights that genetic background drives the splicing patterns of GMAS exons to a similar degree as tissue-specific splicing mechanisms, a previously under-appreciated aspect.

Genes that contain GMAS exons with high tissue variance or high individual variance have substantially different function (Fig. 1B). The first group of genes is enriched in Gene Ontology (GO) terms associated with biophysical properties of the cells, especially related to heart or skeletal muscle function (e.g., sarcomere organization, cardiac muscle development and cytoskeleton organization). This finding supports that alternative splicing is an important aspect contributing to the vast spectrum of biophysical properties of different cell types^30^. In contrast, genes harboring GMAS exons with large variability across individuals are often involved in immune response and signaling pathways. This observation suggests that the individual variability in immune or stress response^31,32,33^ is partly accounted for by splicing variations driven by genetic backgrounds. For genes with GMAS exons with low variability across both tissues and individuals, the most significant GO terms are related to essential cellular processes (Supplemental Fig. S3), which may reflect existence of selection against splicing variability in essential genes.

### GMAS patterns are shared between tissues or individuals

To better understand the tissue-specificity of GMAS events, we next investigated the extent of overlap of GMAS exons between tissues (Methods). We observed that biologically related tissues, such as brain regions, heart and skeletal muscles, and reproductive tissues (uterus and vagina), formed clear clusters (Fig. 2A). Most brain regions shared about 25-43% of GMAS exons with one another, with the exception of cerebellum and cerebellar hemisphere. These two regions were previously reported as outliers with distinctly higher splicing factor expression than other brain regions^29^. Consistently, we observed that these two brain regions shared the most GMAS exons with each other, and much less with other regions (Supplemental Fig. S4).

**Figure 2:**
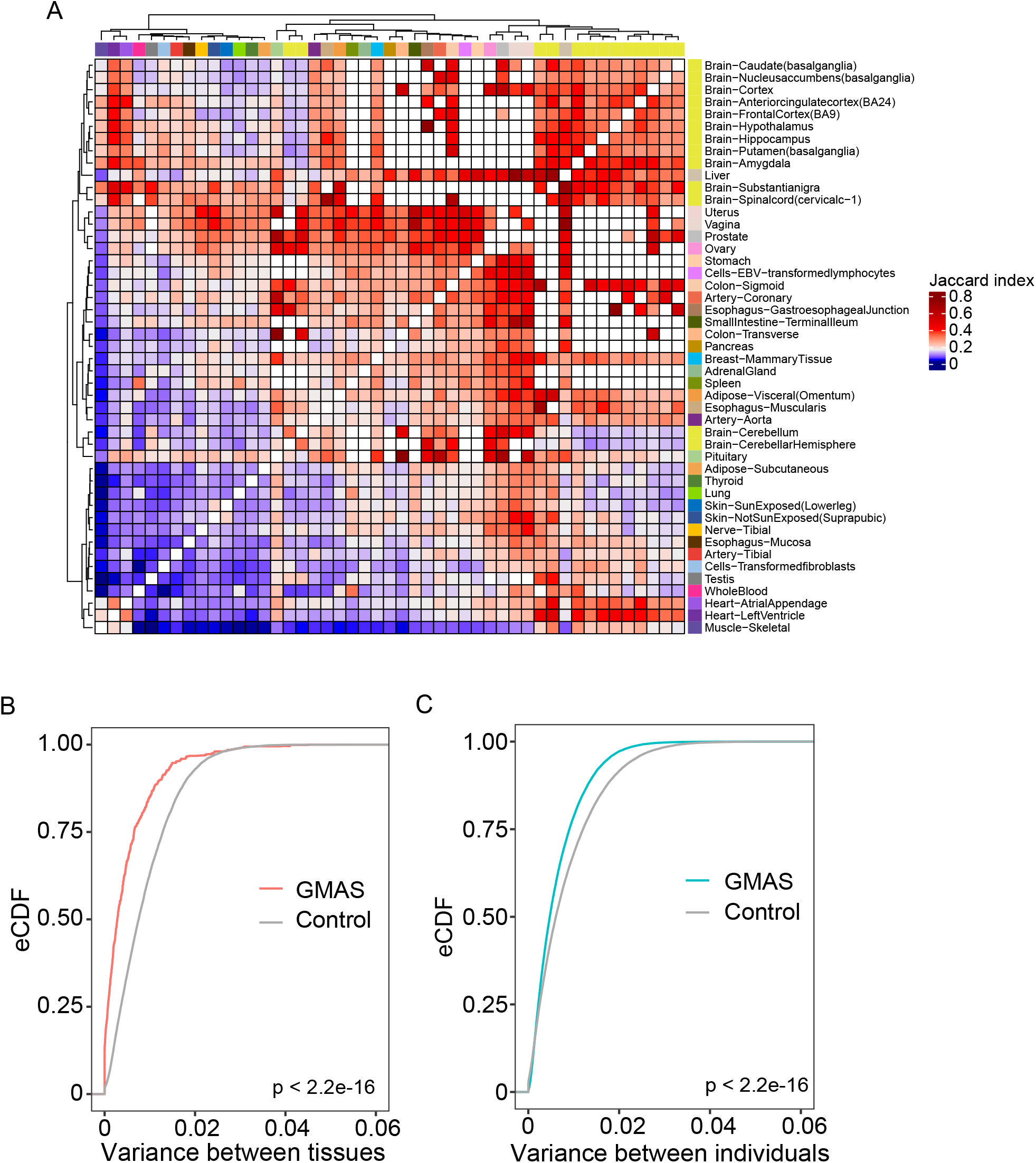
Comparison of GMAS patterns across tissues or individuals. (A) Heatmap of the Jaccard indices of GMAS exons between each pair of tissues (Methods). White boxes correspond to tissue pairs with < 10 common testable exons. (B) Empirical cumulative distribution function (eCDF) of variances across tissues in the allelic biases of tag SNPs of GMAS exons for all individuals. Controls were included for comparison purposes (Methods). The *p* value was calculated using the Kolmogorov– Smirnov test. (C) Similar as (B), but for variance across individuals per tissue.

Next, we asked whether GMAS patterns are shared between distinct tissues more than expected by chance. For this analysis, we focused on 26 representative tissues to remove redundant ones that are highly similar to each other (Methods). Each exon was required to be testable in at least 10 individuals and 5 representative tissues of a specific individual. We observed that the allelic bias of the GMAS tag SNPs was more similar between tissues of the same individual than expected by chance (Fig. 2B). Similarly, for the same tissue, the GMAS-associated allelic bias is also shared among individuals to a greater extent than expected by chance (Fig. 2C).

These results suggest that genetic variants are important drivers for GMAS patterns and tissuespecific effects may play a relatively less dominant role. This observation is consistent with the data in Fig. 1B where the majority of GMAS exons showed relatively small variability across tissues or individuals, with those that are tissue- or individual-specific being the minority.

### Inferring functional SNPs for GMAS events

Since genetic background is a main driver for GMAS events, an important task is to pinpoint the functional genetic variants underlying these events. Note that the tag SNPs identified with the GMAS events are not necessarily functional as they could be in LD with the functional SNPs. To infer the functional SNPs, we developed a new method that combines allele-specific analysis of one dataset with population-level variation in GMAS patterns, namely, concordance-based analysis of GMAS (cGMAS).

Leveraging the genetically driven nature of GMAS, cGMAS is built upon the rationale that a functional SNP, if exists as a heterozygous SNP, should always lead to allele-specific splicing pattern (i.e., GMAS) in the corresponding dataset. Thus, we expect to observe concordance between the genotype of a functional SNP and the splicing patterns of a GMAS exon across different individuals. As illustrated in Fig. 3A (details in Methods), the cGMAS method considers as candidate functional SNPs all heterozygous SNPs in GTEx individuals located in the proximity of GMAS events. For each candidate SNP, a concordance score (*S_i_*) was calculated between its genotype and the GMAS pattern in each individual where the SNP genotype is available. In particular, the GMAS pattern was represented by the allelic imbalance at the tag SNP initially identified with the GMAS event (Supplemental Fig. S5). Subsequently, the distribution of *S_i_* over all individuals was analyzed using a Gaussian Mixture Model (GMM) to identify prominent peaks. The significance of each peak was evaluated via randomization of the *S_i_* values. The functionality of the candidate SNP was determined based on the number and *S_i_* values of significant peaks (FDR ≤ 0.05) detected in the above procedure (Supplemental Fig. S5).

**Figure 3:**
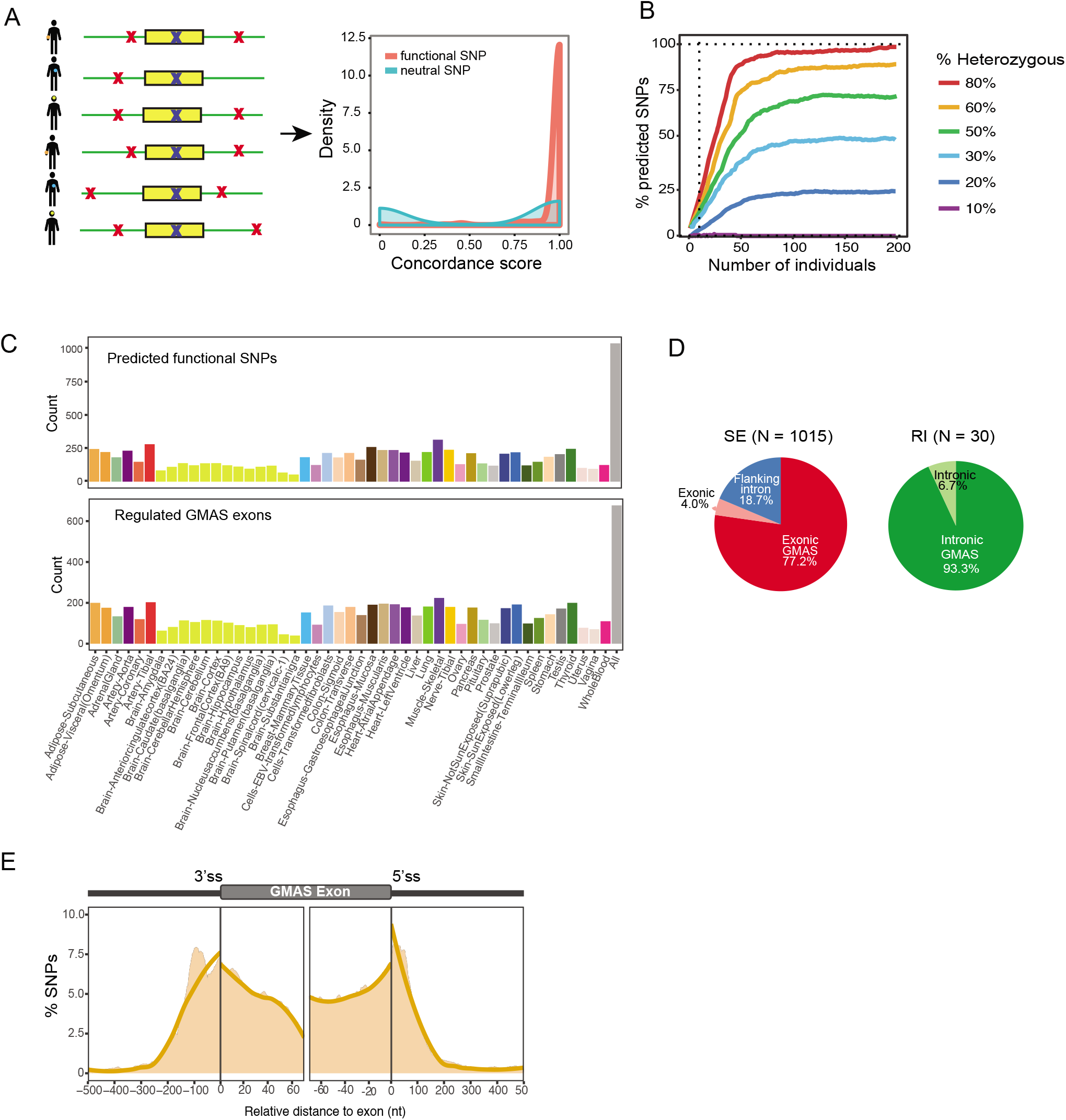
Prediction of functional SNPs for GMAS events. (A) Functional SNPs are predicted by considering candidate SNPs (red crosses) in the vicinity of an GMAS exon, including the tag SNP itself (blue crosses). Concordance among the allelic ratios of the tag SNP in all samples is calculated as described in Methods (with hypothetical distributions shown). (B) The percentage of SNPs predicted given the number of individuals in the simulated testing cohort (Methods). Different percentages of individuals with the heterozygous genotype were simulated. Vertical dotted line marks 10 individuals. (C) Top: number of predicted functional SNPs per tissue. Bottom: number of GMAS exons with predicted functional SNPs per tissue. The rightmost bar (All) corresponds to predictions made by pooling samples from all tissues. (D) Left pie chart: predicted functional SNPs in the exonic or intronic regions of SE (skipped exons). Exonic GMAS: the functional SNP is also the exonic GMAS tag SNP. The rest of the functional SNPs were classified into the “Exonic” or “Flanking intron” group. Right pie chart: for retained introns (RI). Intronic GMAS: the functional SNP is also the GMAS tag SNP. N’s refer to the number of functional SNPs for each group. No functional SNPs were predicted for alternative 5’ or 3’ss exons. (E) Densities of predicted functional SNPs near the exon-intron boundaries of their associated GMAS exons (SEs only). The number of functional SNPs was normalized by the total number of testable SNPs at each nucleotide position. Orange curve is the fitted trend line of the shaded area that represents the SNP density at single nucleotide resolution.

The ability to identify functional SNPs via cGMAS is expected to depend on the number of individuals that possess the GMAS pattern of a given exon. To carry out a power analysis for this method, we simulated 100 hypothetical GMAS exons with functional SNPs that occur in a varying number of individuals (Methods). In addition, we varied the fraction of the simulated individuals that harbor a heterozygous genotype at each functional SNP (Methods). As expected, greater predictive power was achieved if more individuals had the GMAS event (Fig. 3B). The proportion of individuals that had heterozygous alleles at the candidate SNP (i.e., heterozygous ratio) also affected power, where higher heterozygous ratios led to increased power.

### Functional SNPs for GMAS events in GTEx individuals

We applied cGMAS to analyze the GTEx data in two ways: separately for individual tissues and collectively using data of all tissues. Since the number of datasets from each tissue is limited, the latter analysis is associated with increased predictive power. Although tissue-specific functional SNPs may not be identifiable, the pooled analysis could detect SNPs that function relatively ubiquitously across tissues. These analyses together identified 1,045 putative functional SNPs corresponding to 677 GMAS exons (FDR ≤ 0.05, Fig. 3C). These SNPs had 16-24% of overlap with known sQTLs, depending on the method and dataset used for sQTL analyses^22,34,35^ (Supplemental Fig. S6).

Among the putative functional SNPs, about 78% (812) coincided with the GMAS tag SNPs. The rest of the SNPs were located within the same exons as the GMAS exon or in the flanking introns (Fig. 3D). In addition, 23 (2.2%) putative functional SNPs resided in the 5’ splice sites (5’ss), and 26 (2.5%) in the 3’ss. The alternative alleles of these SNPs caused significant difference in the splice site strength (Supplemental Fig. S7). In general, putative functional SNPs demonstrated a positional bias towards enrichment near the splice sites of skipped exons (Fig. 3E), consistent with the expectation that regulatory elements of splicing tend to locate in close proximity to splice sites. Note that since the other types of alternative exons had relatively small numbers of events, they were not included in this analysis.

### Experimental support of functional SNPs for GMAS

To support the predicted functional SNPs, we performed minigene reporter experiments using a splicing reporter from a previous study^36^. For each candidate SNP, we created two versions of the minigene construct, harboring the reference and variant alleles respectively (Supplemental Table 2). Once expressed in cells, minigenes containing functional SNPs are expected to show a significant splicing difference between the two versions. Using this system in HeLa cells, we tested five predicted functional SNPs, three associated with exon skipping events (*PDE4DIP, MAP2K3*, and *UGT2B17*) and two with intron retention events (*SEPT4* and *ATHL1*). All five SNPs were confirmed to lead to allele-specific splicing patterns (Fig. 4A). These results strongly support the predicted functionality of these SNPs.

**Figure 4:**
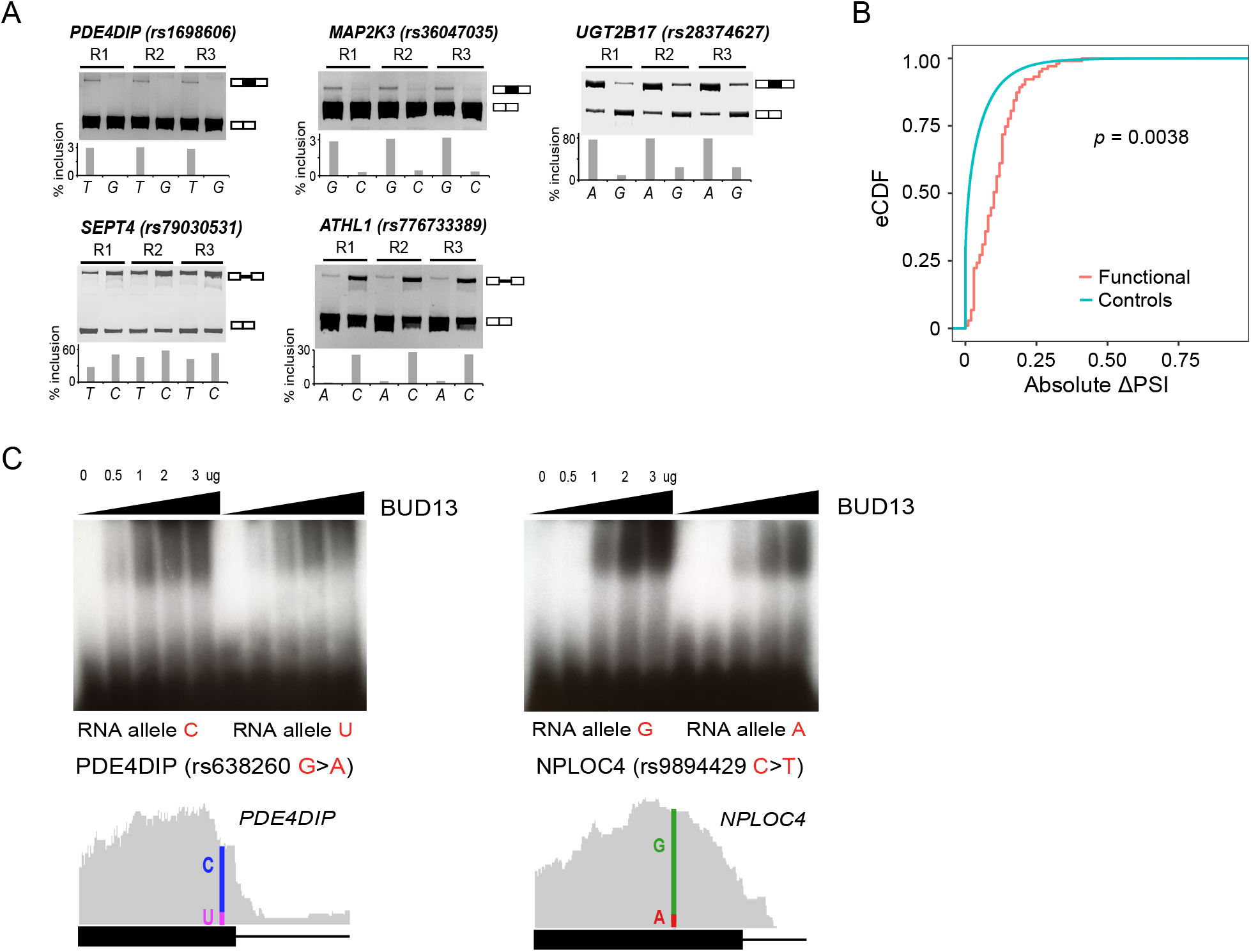
Experimental support of predicted functional SNPs. (A) Minigene experiments validating predicted functional SNPs for GMAS in triplicates (R1-3). The inclusion levels (% inclusion) of the skipped exons or retained introns were estimated from the band intensities of the PAGE gel. (B) eCDF of the absolute changes in the PSI values of GMAS exons upon KD of splicing factors associated with putative functional SNPs (based on motif analysis of eCLIP overlap). The *p* value was calculated using the Kolmogorov–Smirnov test. (C) EMSA validating allele-specific binding of BUD13 to putative functional SNPs. The amount of BUD13 protein used in the experiment is illustrated above the gel. BUD13 eCLIP reads (gray) are shown below the gel, where the locations of the SNPs are labeled with vertical colored lines. The fraction of reads supporting either allele at the functional SNP position is delineated.

It is expected that many functional SNPs may disrupt the interaction between splicing factors and their *cis*-regulatory motifs^6^. Among the putative functional SNPs, 492 were predicted to alter the binding motifs of known splicing factors^37,38^ (using our previous motif analysis method^24^) or overlap the binding sites of splicing factors in the ENCODE eCLIP datasets^39^ (Supplemental Fig. S8A). For these SNPs, we observed that the splicing of their associated GMAS exons showed significant changes upon splicing factor knockdown (KD) compared to random control exons (Fig. 4B), supporting the functional roles of the splicing factors.

Furthermore, 31 putative functional SNPs were testable for allele-specific binding (ASB) using the ENCODE eCLIP data in our previous study^40^, 18 (58%) of which had significant ASB supporting their functional roles. To experimentally confirm the ASB patterns, we carried out electrophoretic mobility shift assays (EMSA, or gel shift) for BUD13, the protein with the largest number of eCLIP peaks overlapping putative functional SNPs (Supplementary Fig. S8A, B). We focused on two candidate functional SNPs and asked whether BUD13 binds to the alternative alleles with different strength. Two versions of the RNA sequences were synthesized harboring the alternative alleles of each SNP. As shown in Fig. 4C, the binding of BUD13 to target RNAs was stronger with increasing protein input. The alternative alleles of the SNPs demonstrated visible differences in their binding to the protein, supporting the functional impact of these SNPs. Note that the two SNPs were also predicted as ASB SNPs for BUD13 via eCLIP-seq analysis of K562 cells^40^ (Fig. 4C), consistent with our experimental results.

### GMAS events are enriched in disease-relevant genes and regions

To examine the disease relevance of GMAS events, we first asked whether GMAS events are significantly associated with GWAS loci. For this analysis, we included all GMAS events identified in this study, not limited to those with predicted functional SNPs. Specifically, we examined whether GMAS-associated SNPs were in LD with GWAS SNPs (and within 200kb in distance, Methods). As controls, random variants from non-GMAS genes were sampled and analyzed relative to GWAS SNPs similarly. We observed that 72% (5317) GMAS SNPs were in LD with GWAS SNPs, a percentage significantly higher than that among control SNPs (Fig. 5A; P < 2.2 × 10^−16^). Note that similar results were observed when only including the putative functional SNPs for GMAS (Fig. 5A; P < 2.2 × 10^−16^). These observations support the likely disease relevance of GMAS events.

**Figure 5:**
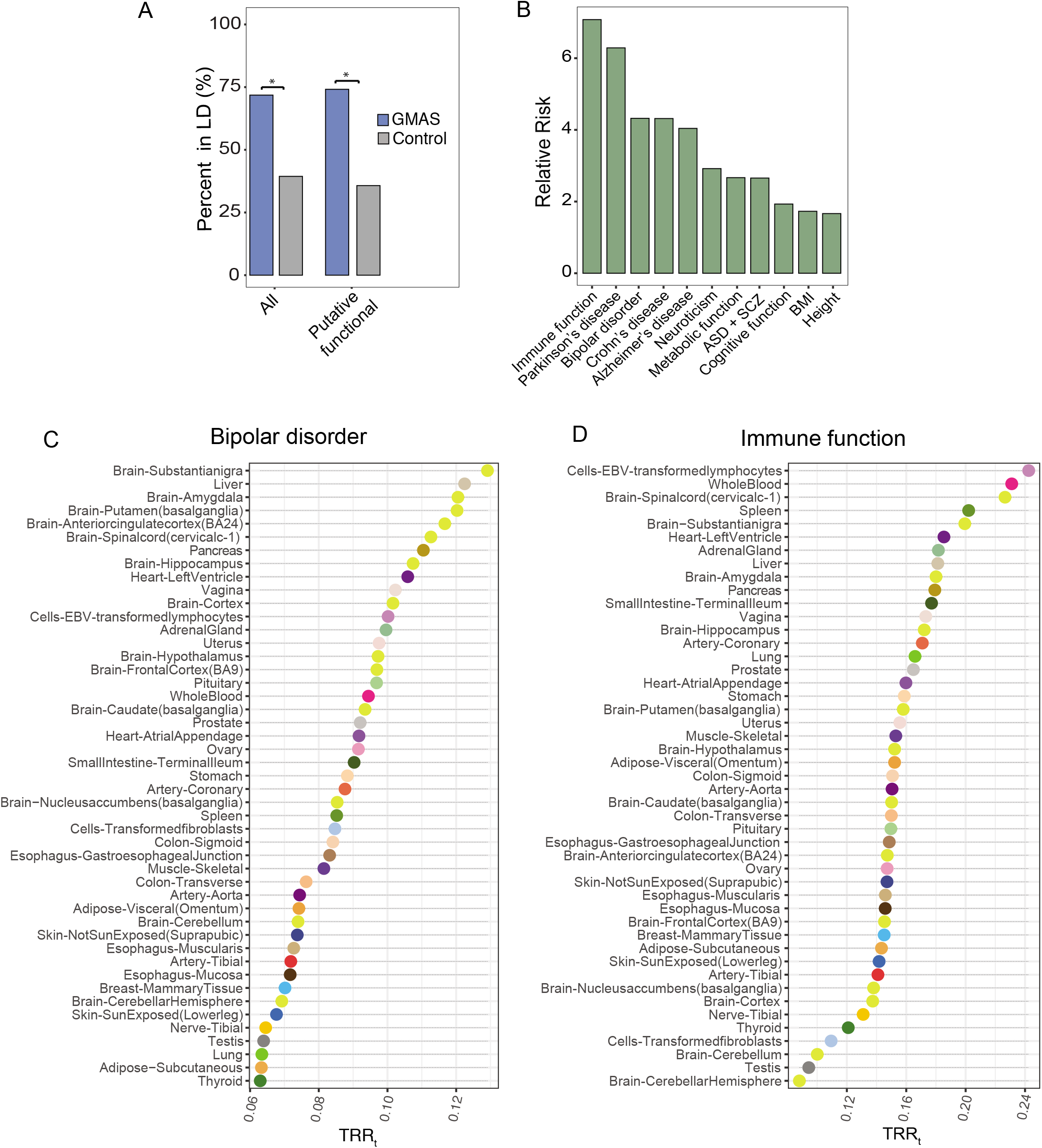
Functional relevance of GMAS events. (A) Proportions of all GMAS SNPs and putative functional SNPs in LD with (and within 200kb of) GWAS SNPs. Controls were random dbSNPs in genes that do not harbor GMAS SNPs. *\*p* < 2.2 x 10^−16^ (Fisher’s Exact test) (B) Relative risk of GMAS SNPs in LD with selected trait/disease, defined as the ratio between the values in (A) of the GMAS and control groups for all GMAS SNPs. ASD: autism spectrum disorder. SCZ: schizophrenia. BMI: body mass index. All p values < 2.2 x 10^−16^ (Fisher’s Exact test). (C)-(D) Trait-relevance ratio of each tissue (TRR_t_) defined as the proportion of GMAS SNPs identified in each tissue that were in LD with GWAS SNPs, for bipolar disorder and immune function, respectively.

We further examined the GMAS-GWAS relationship for specific traits/diseases. For each trait/disease, we repeated the above LD analysis and calculated the enrichment of GMAS SNPs relative to control SNPs that are located in LD regions of GWAS SNPs (defined as relative risk, Methods, Fig. 5B). A number of traits/diseases, such as immune function, Parkinson’s disease, and Bipolar disorder, demonstrated significantly high enrichment (Fig. 5B). Interestingly, GMAS SNPs associated with immune function had the highest relative enrichment, consistent with the known prevalence of alternative splicing in the immune system^41^. The enriched association of GMAS SNPs with neurological function or related diseases suggests that splicing may have close relevance in these processes. Complex traits such as height and body mass index (BMI) had the lowest relative risk (although still significant), indicating that splicing likely contributes the least to their underlying biological mechanisms among those considered here.

An interesting question is whether the GMAS events were identified in tissues relevant to their associated GWAS traits/diseases. To this end, we defined a trait-relevance ratio (TRR) to evaluate the proportion of GMAS SNPs in each tissue that were in LD with GWAS SNPs for a given trait/disease (Methods). This analysis revealed some interesting insights. For example, for Bipolar disorder, brain tissues had the highest TRRs among all tissues, consistent with the nature of the disease (Fig. 5C). In contrast, TRRs were highest in lymphocytes and whole blood for immune function-associated GMAS SNPs (Fig. 5D), both with immune relevance. In addition, GMAS SNPs associated with metabolic function had highest TRRs in tissues (liver and adrenal gland) of close relevance to metabolism (Supplemental Fig. S9A). Neuroticism- and cognitive function-related GMAS SNPs were observed with high TRRs in brain tissues (Supplemental Fig. S9B, S9C). Thus, these observations are highly consistent with the expected tissues of relevance of the traits/diseases, supporting the potential involvement of GMAS in related functional processes. For other traits, the top tissues with high TRR values were more diverse or non-intuitive (Supplemental Fig. S9D-S9I). It’s likely that genetically driven splicing alteration is not a primary contributor, or alternatively, these traits/diseases are complex and involve biological processes in a wide range of tissues.

## Discussion

We report a comprehensive study of allele-specific alternative splicing (a.k.a. GMAS) in human tissues. Using GTEx datasets, we identified thousands of GMAS events, encompassing 4941 exons and 7404 SNPs. The multifaceted nature of the data allowed an examination of the GMAS landscape across tissues and individuals. We observed that the allele-specific pattern of GMAS events varied to similar degrees across tissues and individuals. It is well-established that alternative splicing demonstrates high tissue-specificity, which enables segregation of samples by tissue types rather than per individual^29,42,43^. In contrast, our analysis showed that, for genetically regulated splicing events, the genetic contribution to splicing variability is equivalent to that contributed by tissue-specificity. As tissue-specificity is often imposed by *trans*-acting regulators, our results suggest that *cis*- and *trans*-regulatory mechanisms have similar degrees of impact on the variability of GMAS.

In general, GMAS events can be shared across tissues or individuals, or demonstrate high tissue- or individual-specificity (Fig. 1B, Fig. 2). We observed that GMAS events overall are shared more significantly than expected by chance across tissues or individuals (that share the same genotype) (Fig. 2B, C). This result is consistent with previous literature that genetically driven splicing profiles tend to be common to different cell or tissue types^22,23,24^. This is expected because genetic determinants are the most important factor for such splicing events. On the other hand, there do exist many GMAS events that are highly individual- or tissue-specific (Fig. 1B). Interestingly, genes with individual-specific GMAS exons are often involved in immune-related processes. This observation not only highlights the impact of an individual’s genetic makeup on the immune system, but also identifies splicing as a potential mechanism through which the phenotypic effects of genetic variants are manifested. In contrast, genes containing GMAS exons with high tissue variability are involved in heart or skeletal muscle function, supporting the particular importance of alternative splicing in the biophysical properties and functions of cells^44^.

Leveraging the GTEx genotype information and GMAS events, we developed a new method to pinpoint functional SNPs that regulate splicing. Specifically, our method appraises the concordance between the allelic bias of a candidate SNP and the splicing pattern of an alternatively spliced region, as represented by the allelic signature of the tag GMAS SNP. The key factor that determines the performance of our method is the “heterozygous ratio” of a candidate functional SNP among the testing cohort. Our method demonstrates high predictive power when many individuals have heterozygous alleles at the candidate SNP locus. Within the GTEx cohort, we were able to predict over 1000 functional SNPs for GMAS, and the quality of our predictions was confirmed by the enrichment of functional SNPs near the splice sites, a popular metric used to examine the splicing relevance of a SNP. This method can be generally applied to any dataset encompassing large populations to expand the repertoire of functional SNPs that regulate splicing.

Many large-scale efforts have been devoted to understanding the functional relevance of SNPs in the human genome. To date, the GWAS catalog has documented hundreds of thousands of phenotype-associated SNPs from over 3500 publications^2^. Yet, many traits were found to associate with non-coding or intergenic SNPs that do not alter the protein sequences, which makes GWAS interpretation challenging. We observed that a high fraction of GMAS events are associated with SNPs in LD with GWAS loci, suggesting that these GWAS-reported SNP-trait associations may be related to dysregulation of splicing. This observation is further substantiated by the GMAS enrichment in tissues of expected relevance for a number of GWAS traits (e.g., bipolar disorder, metabolic and immune function). Our study indicates that allele-specific splicing analysis is an effective means to discover functionally relevant genetic variants that may contribute to disease mechanisms.

## Methods

### Preprocessing of GTEx RNA-seq data and identification of GMAS events

FASTQ files from individuals with genotype information (from whole genome sequencing, whole exome sequence, or Illumina SNP Arrays) were downloaded from the GTEx database (v6p release). Library adaptors were trimmed by Cutadapt^45^. We aligned the reads to the hg19 genome and transcriptome using HISAT2^25^ with parameters --mp 6,4 --no-softclip --no-mixed --no-discordant, keeping only the uniquely mapped read pairs for the following analyses. Samples with fewer than 25 million uniquely aligned read pairs were considered as insufficient read coverage for detecting GMAS events and thus discarded (about 10% of all datasets). We focused on the tissues that have at least 50 samples with sufficient read coverage. In total, 7822 RNA-seq samples across 47 tissues from 515 donors were kept for the GMAS analysis.

We collected a list of high-quality SNPs from whole genome sequencing (quality filter: GQ ≥ 20), whole exome sequence (quality filter: GQ ≥ 20), and Illumina SNP Arrays (quality filter: IGC ≥ 0.2) provided by GTEx. In addition to the genotyped SNPs, we included all dbSNPs (version 146) that showed RNA-seq evidence of being heterozygous in at least one GTEx individual as potential candidates for the GMAS analysis. To determine which dbSNPs were heterozygous, we used the RNA-seq reads covering the candidate dbSNP position and defined the SNP to be heterozygous if it had at least 3 reads for each of the two alleles (with at least 20 total reads). Additionally, we further filtered out those with extreme allelic ratio (AR, defined as number of reads covering the reference allele/total number of reads), i.e., AR < 0.1 or AR > 0.9, to avoid potential amplification biases or sequencing errors.

We applied our published method^23^ to predict GMAS events using the combined list of SNPs (genotyped or RNA-seq-based) and the uniquely aligned RNA-seq reads. Briefly, this method first examines allele-specific expression (ASE) of all heterozygous SNPs in a gene. It then determines whether ASE is global in the specific gene, which represents gene-level ASE possibly regulated at the level of transcription or RNA decay that affects all heterozygous SNPs in the gene. Alternatively, a gene may have local ASE, that is, ASE demonstrated in only a small fraction of testable (− 20 read coverage) heterozygous SNPs. GMAS accounts for a type of such local ASE patterns, where the ASE SNP is located in an alternatively spliced exon and has significant allelic bias relative to control SNPs in the same gene (non-ASE SNPs).

Relative to the published version^23^, we made the following modifications in this study. First, instead of focusing solely on annotated alternatively spliced exons from GENCODE comprehensive annotation (v24lift37), we tested all internal exons for potential GMAS events. Second, we replaced the normalized expression value (NEV) by PSI calculated by the method described in Schafer et al. 2015^46^, only keeping exons with ≥ 15 total reads (inclusion reads + exclusion reads) or ≥ 2 exclusion reads. An exon is testable if it passes the read coverage requirements for PSI calculation and has a powerful (defined as having ≥ 20 read coverage) non-ASE SNP in another exon of the same gene^23^. To avoid false positives, we only focused on GMAS events that were called in at least three samples out of the total 7822 samples we analyzed.

### Estimation of tissue vs. individual contributions to GMAS pattern variations

We used the lmer function from the lme4 package in R to model the allelic imbalance for each GMAS exon as the following:

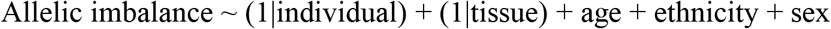

The allelic imbalance was calculated as the absolute difference of allelic ratio to 0.5. The fixed effects (age, ethnicity, and sex) were chosen based on the previous literature^29^. The allelic imbalance variations contributed by tissues and by individuals were estimated from the above model.

### Tissue-specificity quantified by Jaccard index and GMAS frequency

We used the Jaccard index to quantify the extent of sharing of the GMAS pattern for an exon *e* between tissues *i* and *j* (*s_eij_*). Specifically, 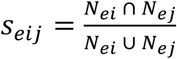, where *N_ei_* and *N_ej_* are the number of individuals with *e* showing GMAS pattern in tissues *i* and *j* respectively (*i* ≠ *j*). To reliably estimate *s_eij_*, we required *N_ei_* ∪ *N_ej_* ≥ 10. The final GMAS pattern shared between tissues *i* and *j* (*s_ij_*) was calculated as 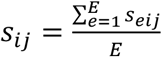, where *E* is the total number of exons with *s_eij_* for tissues *i* and *j*.

### Tissue and individual variability in GMAS

To assess the variability in GMAS across individuals and tissues, we used variance as a quantitative measure of dissimilarity in allelic biases. For each exon showing GMAS pattern in any given individual, we measured the variance within allelic biases of the tag SNPs in all corresponding tissues of the individual. As controls, we sampled allelic biases of the tag SNPs of the same exon in similar tissues but different individuals and calculated their variance. The distribution of variances across all individuals for the GMAS exons was then compared to that of the controls (Fig. 2B). Similarly, for each exon showing GMAS pattern in a given tissue, we calculated the variance among the allelic biases of the tag SNPs across individuals. The controls were randomly sampled allelic biases of the tag SNPs of the same exon in individuals showing GMAS pattern for the exon but different tissues. Again, we compared the distribution of variances across all tissues for the GMAS exons to the distribution of variances in controls (Fig. 2C).

### Prediction of functional SNPs for GMAS

The basic rationale for our method is that a functional SNP for GMAS should show concordant relationship (cGMAS) between its genotype and the splicing pattern of the target exon across a large number of individuals. In the toy example illustrated in Supplemental Fig. S5A, we first define a distance metric *d* = |0.5-*R*_tag_|, where *R*_tag_ is the allelic ratio of the tag SNP defined as *N*_ref_/(*N*_ref_ + *N*_alt_). *N*_ref_ and *N*_alt_ denote the number of reads harboring the reference allele and the alternative allele of the SNP, respectively. Thus, *d* represents the difference between the allelic ratio of the tag SNP and the expected allelic ratio of an unbiased SNP. In Supplemental Fig. S5A, the candidate SNP (which is different from the tag) is assumed to be the functional SNP underlying GMAS, with the A allele causing exon inclusion and G allele causing exon skipping. Thus, for individuals with the homozygous genotype (AA or GG) at the candidate SNP, *d* is expected to be 0.5. On the other hand, for individuals with AG genotype at the candidate SNP, *d* is 0 or 1 depending on the haplotype between the tag and candidate SNPs.

Next, we define the concordance score (*S_i_*) for this example exon in individual *i*, similarly as used in a previous study^35^. *S_i_* measures the concordance level between the genotype and the splicing pattern.

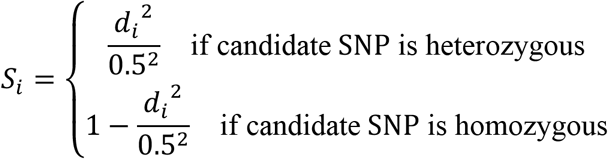

For the toy example in Supplemental Fig. S5A where A/G alleles of the candidate SNP cause complete switch of exon inclusion/exclusion, the value of *S_i_* is 1. In a different scenario as illustrated in Supplemental Fig. S5B where the tag SNP is considered as the candidate functional SNP, we define:

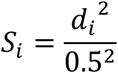

Thus, in case of a functional SNP causing complete switch of exon inclusion/exclusion, the value of *S_i_* is also 1. In general, for true functional SNPs, *S_i_* is expected to have a distribution with a peak close to 1, whereas random neural SNPs have broadly distributed *S_i_* values (Supplemental Fig. S5C).

For more realistic cases where the two alleles of the functional SNP do not cause 100% splicing difference, the distribution of *S_i_* is multi-modal. In addition to a peak close to 1, another peak in the medium *S_i_* range (>0) exists. On the other hand, a peak at 0 corresponds to non-functional SNPs. To unbiasedly model the distribution of *S_i_*, we fitted a Gaussian Mixture Model (GMM) to identify its peaks. The number of GMM components was determined via the Bayesian information criterion (BIC). A z-test was carried out to search for peaks whose average values were significantly different from 0 (FDR ≤ 0.1). For a true functional SNP, the *S_i_* distribution should be supported by individuals with different genotypes (homozygous or heterozygous). To avoid potential false positives driven by a specific genotype in a small number of individuals, we excluded candidate SNPs where the genotype supporting the *S_i_* peak is significantly biased towards one genotype (Fisher’s exact test, FDR ≤ 0.1).

To ensure the magnitude of the peak was significant, we binned the x-axis (*S_i_* scores) into 100 bins and randomized the data points evenly across the bins to generate a background distribution. This process was repeated 500 times to estimate an average background peak level and its standard deviation. We compared the peak height to the background in the same bin and defined significant peaks by z-score > 2.58, which corresponds to *P* < 0.01.

For each GMAS exon, we examined all SNPs in the exon and the immediate introns as candidate functional SNPs (Supplemental Fig. S5D). SNPs that are homozygous in all individuals were not considered. The concordance score for each candidate and tag SNP pair was calculated and the functional SNP was predicted as described above.

### Power analysis for predicting functional SNPs for GMAS

To assess how many individuals our method necessitates to predict functional SNPs for GMAS, we simulated 100 functional SNPs with two alternating alleles inducing 75% difference in PSI. This allele-specific splicing difference is reflected in the allelic ratios. The total read counts of a SNP were simulated from a negative binomial distribution using parameters estimated from a real GTEx RNA-seq sample. We required all simulated SNPs to have at least 20 reads. The allelic ratios of the simulated SNPs were generated from a binomial distribution.

We simulated six groups of 200 individuals. Each group has a specific heterozygous frequency (Fig. 3B), which is defined as the fraction of individuals with heterozygous alleles at the candidate SNP position in a group. We ran the cGMAS method on the 100 SNPs by varying the number of individuals while maintaining the heterozygous frequency for prediction. Figure 3B illustrates the power of this method in the different simulations.

### Analysis of ENCODE eCLIP-seq and RNA-seq data

eCLIP peaks were obtained from the ENCODE portal (https://www.encodeproject.org). The ENCODE RNA-seq data were analyzed similarly as described above for GTEx RNA-seq data. PSI values of replicated samples were averaged in Fig. 4C.

### Analysis of GMAS SNPs in LD with GWAS SNPs

Trait-variant associations with p-values larger than 5.0 × 10^−8^ were removed from the GWAS catalog^2^ (version 1.0.2 – downloaded 2020-02-04). In addition, the GWAS SNPs were separated into LD blocks according to the LD information of the CEU population and further required to have R^2^ ≥ 0.8 and D’ ≥ 0.9. To evaluate the functional relevance of GMAS SNPs with regard to GWAS, we calculated the number of GMAS SNPs in LD with and within 200kb of at least one GWAS SNP (referred to as GMAS-GWAS SNPs). A similar number was also calculated for the putative functional SNPs. To determine the significance of the above enrichment, we randomly sampled the same number of dbSNPs from genes that do not host GMAS events. The number of randomized dbSNPs in LD with and within 200kb of at least one GWAS SNP was compared to that of the GMAS SNPs with a Fisher’s Exact test.

To investigate the enrichment of GMAS-GWAS SNPs in specific traits/diseases, we calculated the relative risk of GMAS SNPs being in LD with and within 200kb of a GWAS SNP for the trait of interest versus control SNPs. The relative risk or risk ratio was calculated as follows:

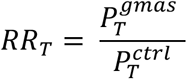

where *RR_T_* is the relative risk for trait 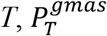 is the proportion of GMAS SNPs in LD with GWAS SNPs for trait *T* and 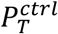 is the proportion of control SNPs in LD with GWAS traits for trait *T*.

As a measure of how relevant the GMAS-GWAS SNPs are to the corresponding traits, we calculated the trait-relevance ratio (TRR_t_) for each tissue in which the SNP showed GMAS pattern. The TRRt metric controls for the number of GMAS events identified per tissue and is calculated as:

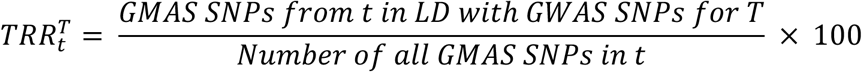

where 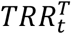 is the trait-relevance ratio, *T* is the trait of interest and *t* is a source tissue of a GMAS-GWAS SNP.

### Cell culture

HEK 293T and HeLa cells were obtained from ATCC and maintained in DMEM supplemented with 10% FBS (ThermoFisher Scientific, 10082147) and antibiotics at 37°C in 5% CO_2_.

### Construction of minigenes

Minigenes containing SNP candidates were cloned as previously described^40^. Briefly, the candidate skipped exon and ~500 nt of each flanking intron were amplified using HeLa genomic DNA. The DNA fragment was then sub-cloned into the pZW1 splicing reporter using HindIII and SacII or EcoRI and SacII cloning sites. The candidates for intron retention were cloned into pcDNA3.1 plasmid. Final constructs were confirmed by Sanger sequencing. Primers used in this study are listed in Supplemental Table 2.

### Transfection, RNA extraction, reverse transcription, and PCR

Minigene constructs were transfected into >90% confluence HeLa cells using Lipofectamine 3000 (ThermoFisher Scientific, L300015). Total RNA was isolated after 24 h transfection using TRIzol (ThermoFisher Scientific, 15596018) followed by Direct-zol RNA Miniprep plus kit (Zymo Research, R2072). cDNA was produced from 2 μg of total RNA by SuperScript IV First-Strand Synthesis System (ThermoFisher Scientific, 18091050). To amplify the candidate exons in minigene constructs, 5% of the cDNA was used as template via 26 PCR cycles (Supplemental Table 2).

### Gel electrophoresis and quantification

The PCR amplicon was loaded onto 5% polyacrylamide gel and ran at 70 volts for 1.5 hours. The PAGE gel was stained with SYBR^®^ Safe DNA Gel Stain (ThermoFisher Scientific, S33102) for 30 min and the gel image was taken by Syngene SYBRsafe program (Syngene). Spliced isoforms expression level was estimated using the ImageJ software (http://imagej.nih.gov/ij/). Inclusion or intron retention rate (% inclusion) of the target exon was calculated as the intensity ratio of upper/(upper+lower) bands.

### Cloning of human BUD13 and lentiviral overexpression

Human BUD13 was cloned from HeLa cDNA into the pCR 2.1-TOPO vector (Thermo Fisher Scientific, 450641). After sequence confirmation, BUD13 was sub-cloned into the pcDNA3.1 backbone containing 3×Flag-6HIS tag using NotI and EcoRI sites. To achieve stable overexpression, the 3×Flag-BUD13-6HIS fragment was transferred into the pLJM1 lentiviral construct using the NdeI and EcoRI sites (Addgene plasmid # 19319). We produced lentiviruses via co-transfection of pCMV-d8.91, pVSV-G and pLJM1-3×Flag-BUD13-6HIS into HEK293T cells using Lipofectamine 3000 (Thermo Fisher Scientific, L3000015). Lentiviruses were collected from conditioned media after 48 h co-transfection and filtered through 0.2μm syringe filter. Lentivirus-containing medium was mixed with the same volume of DMEM containing polybrene (8μg/mL). The lentiviruses were transduced into HEK293T cells in ten 150mm culture plates, where they were incubated with 2μg /mL puromycin for 48h.

### Purification of recombinant human BUD13

HEK293T cells stably expressing BUD13 were centrifuged at 1000 × g for 5 min at 4°C and the pellets were resuspended with ice-cold 5mL lysis buffer (PBS, 20mM Imidazole, 0.5% IGEPAL CA-630, 0.5mM DTT, 0.5 × protease inhibitor cocktail, 100U DNAse I). After 30 min incubation, the lysate was disrupted using sonication at 25% amplitude for 20sec with 1sec pulse. Next the lysate was centrifuged at 13,000 × g for 5 min at 4°C. The supernatant was collected and filtered using 0.45μm syringe filter. The sample was incubated with 1mL Ni-NTA agarose (Thermo Fisher Scientific, R90110) for 6 hrs at 4°C followed by five times of washing with 5mL buffer A (PBS, 20mM Imidazole, 0.5mM DTT, 0.5% IGEPAL CA-630, 0.5 × protease inhibitor cocktail). Proteins were eluted with 3mL elution buffer (PBS, 250mM Imidazole) and excess salt was removed using the desalting column according to the manufacture’s protocol (GE Healthcare, 17085101). Subsequently, Flag affinity purification was performed using 1mL Flag agarose bead (MilliporeSigma, A2220) according to the manufacture’s protocol. Elution was performed using 100mg/mL counter flag peptide. Flag peptide and small size of non-specific proteins were removed by 20K Slide-A-Lyzer dialysis cassette (ThermoFisher Scientific, 66003) with 1L binding buffer (PBS, 0.5% IGEPAL CA-630, 5% glycerol) in the cold room overnight. Recombinant BUD13 purification was confirmed by SimplyBue SafeStain (Thermo Fisher Scientific, LC6060) and western blot using BUD13 antibody (Bethyl Laboratories, A303-320A). Protein concentration was measured by Pierce Coomassie (Bradford) protein assay kit (ThermoFisher Scientific, 23200) and the Turner spectrophotometer SP-830.

### In vitro transcription of BUD13 target RNA

Sense and antisense oligos including T7 promoter (Supplemental Table 2) were annealed at 95°C for 5min in a heat block then cooled down to room temperature for 3 hrs. *In vitro* transcription was performed using HiScribe T7 high yield RNA synthesis kit according to the manufacturer’s protocol (NEB, E2040S). *In vitro* synthesized RNAs was treated with 10U RNAse-free DNAse I (ThermoFisher Scientific, EN0525) at room temperature for 30min, then purified by the RNA clean & concentrator-5 Kit (Zymo Research, R1015). Next, RNA samples were treated with 10U shrimp alkaline phosphatase (NEB, M0371S) at 37°C for 1 hr and then labeled with 0.5μl of gamma ^32^P-ATP (PerkinElmer, BLU502A250UC) using 20U T4 polynucleotide kinase (NEB, M0201S). Subsequently RNA probes were purified by 5% Urea PAGE extraction and RNA clean & concentrator-5 Kit. RNA concentration was measured by Qubit 2.0 fluorometer (ThermoFisher Scientific).

### Electrophoretic Mobility Shift Assay (EMSA)

The RNA probes and recombinant BUD13 protein (0, 0.5, 1, 2, and 3 μg) were incubated in 15μl of the binding buffer (PBS, 0.5% IGEPAL CA-630, 5% glycerol, 0.1 × protease inhibitor cocktail, 10U RNAse inhibitor) at 28°C for 30 min, then loaded onto 5% TBE-PAGE run at 75V for 1.5 hrs. The gel was processed without drying, covered with clear folder and exposed to X-ray film at −80°C.

## Acknowledgements

We thank members of the Xiao laboratory for helpful discussions and comments on this work. We thank the GTEx Consortium for generating the valuable datasets used in this study. We thank the ENCODE Consortium (Graveley and Yeo groups) for generating the eCLIP and RNA-seq data. This work was supported in part by grants U01HG009417 and R01AG056476 to XX. KA was supported by the UC-HBCU Initiative Fellowship from the University of California Office of the President. YHH was supported by the Bioengineering supplemental fellowship of UCLA.

