## Supplemental Figures for "Allele-specific alternative splicing in human tissues"

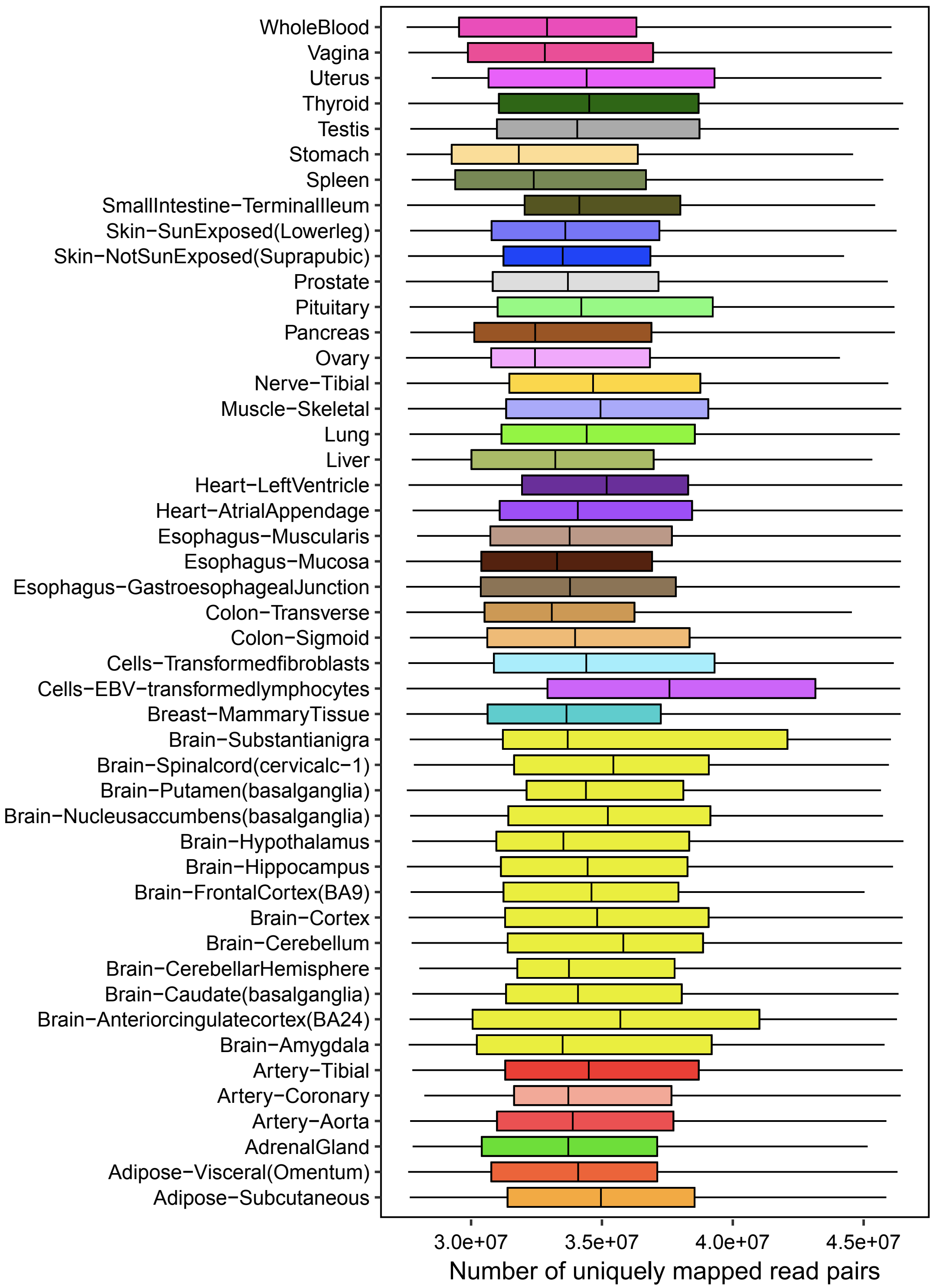


Figure S1. Number of uniquely mapped read pairs per tissue.

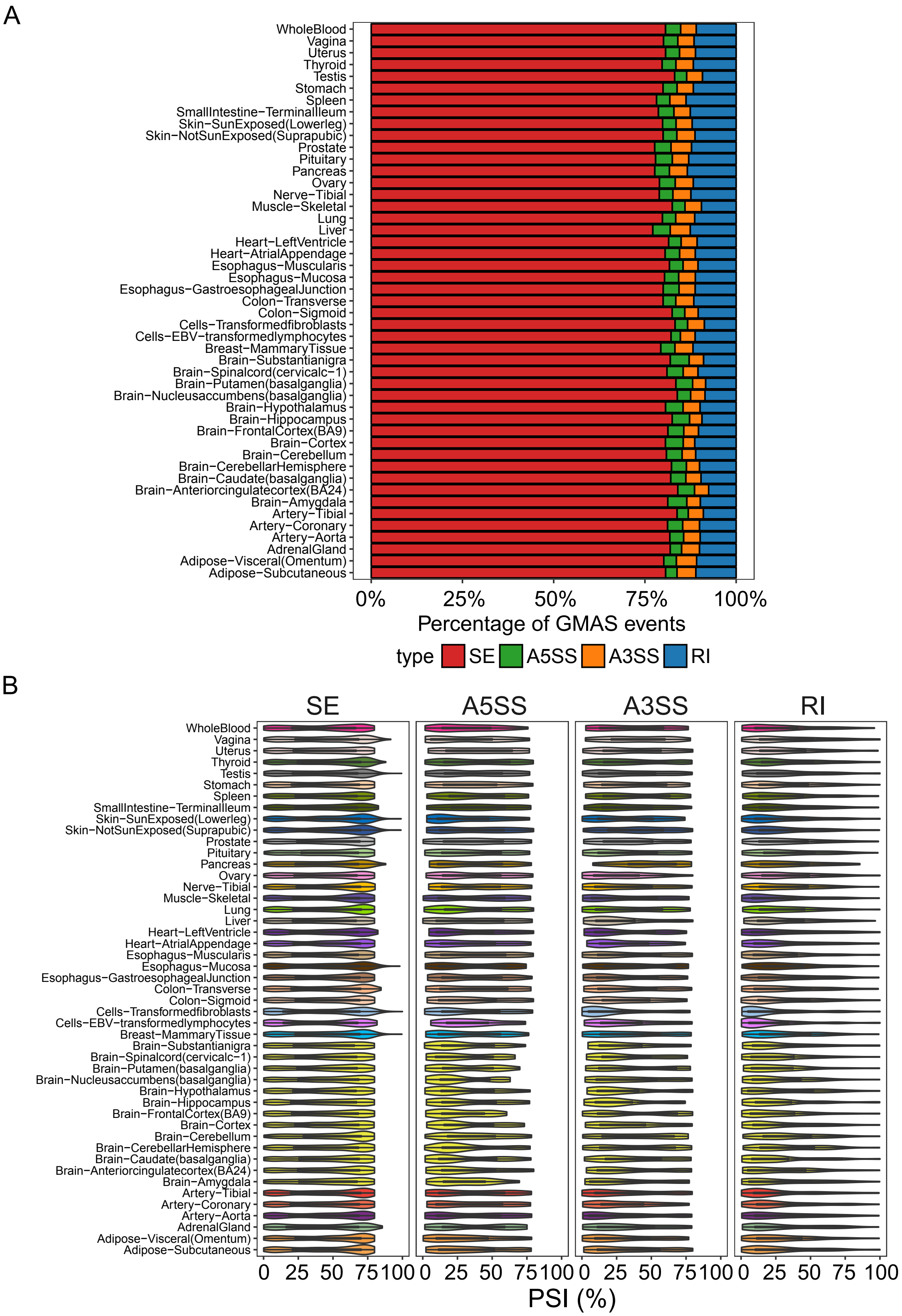


Figure S2. (A) Distribution of GMAS types per tissue. (B) The PSI distribution of GMAS exons per tissue of 4 different types of splicing events. SE: skipped exon; A5SS: alternative 5’ss exon; A3SS: alternative 3’ss exon; RI: retained intron.

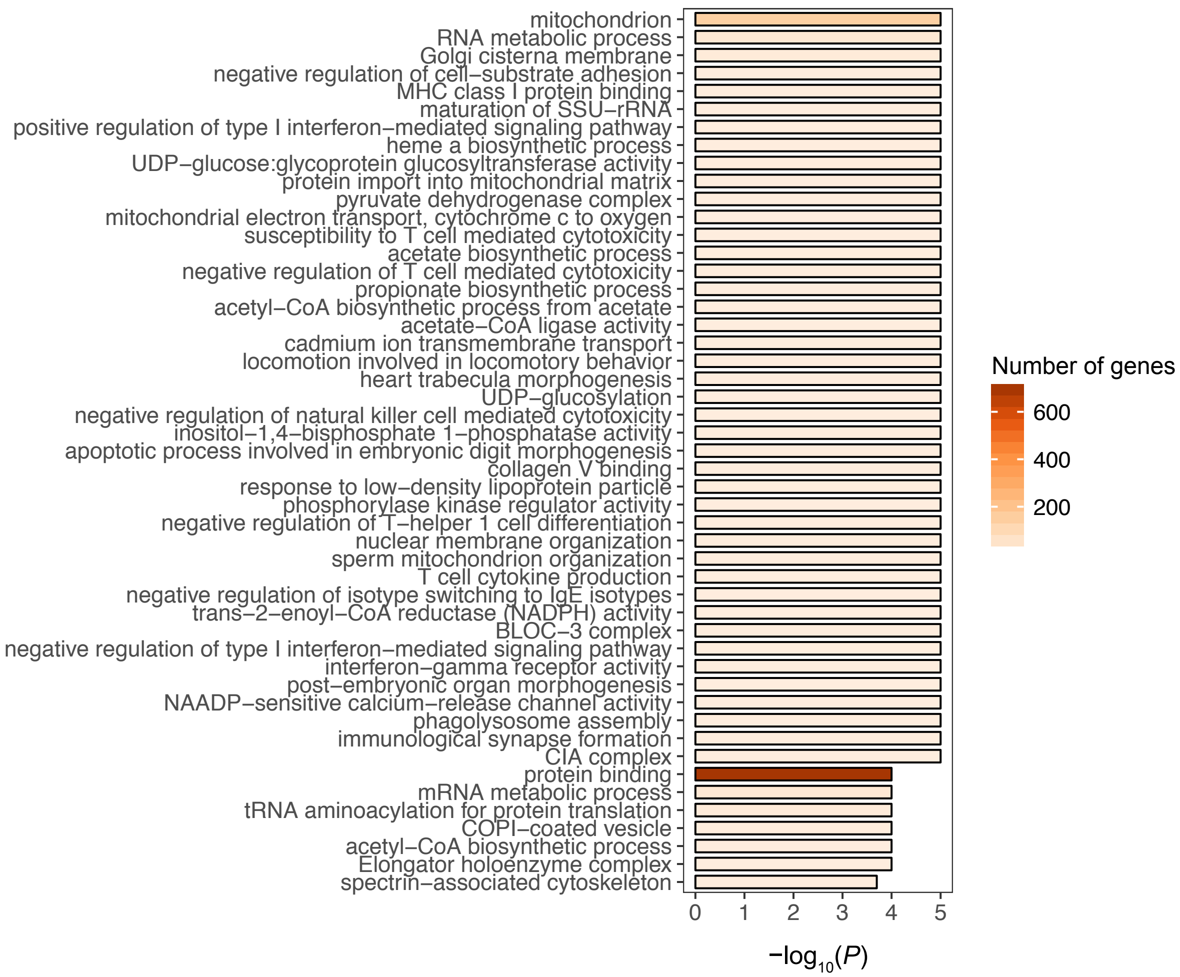


Figure S3. GO terms enriched among genes harboring GMAS exons with low variability across tissues and across individuals (N = 1636).

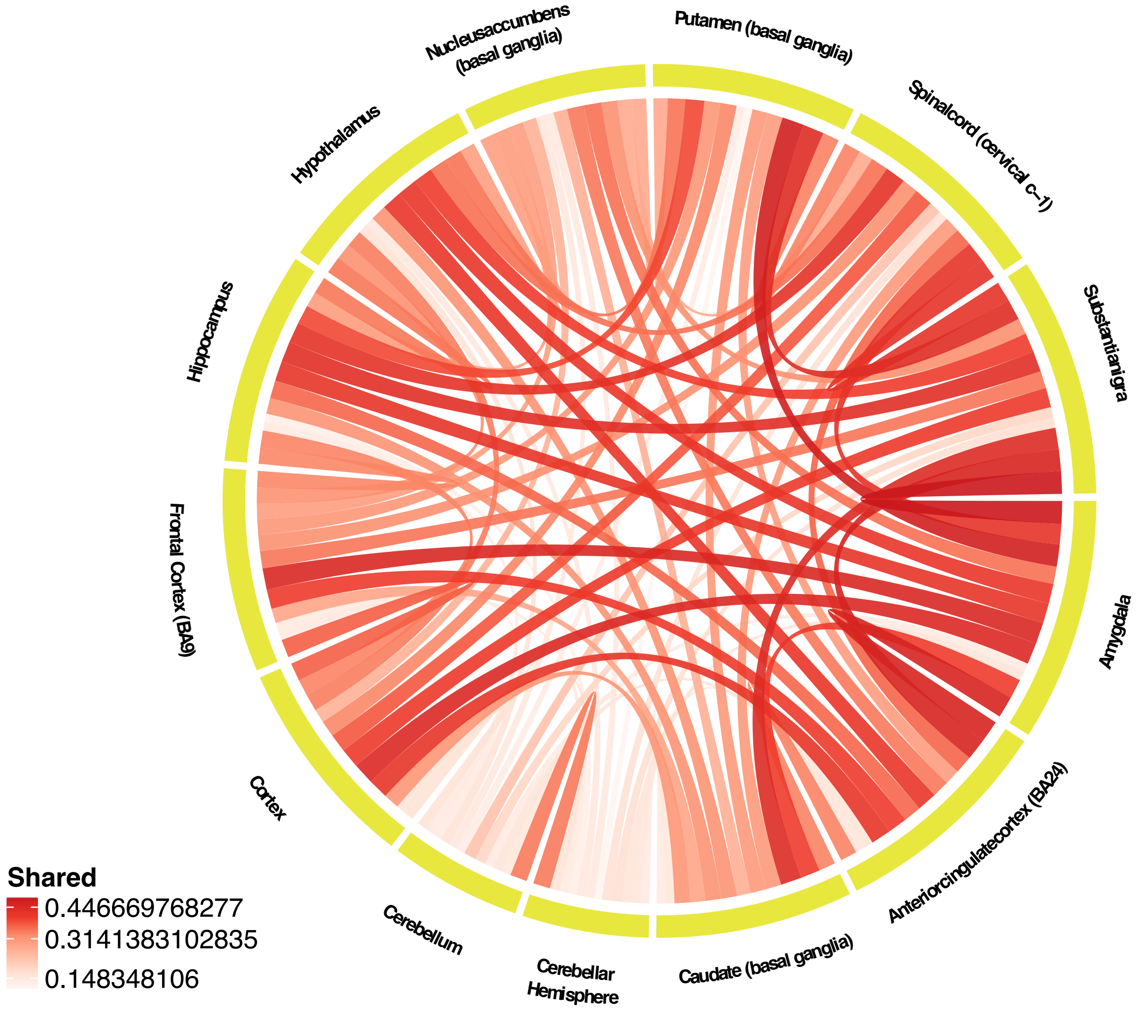


Figure S4. Jaccard index of GMAS patterns between pairs of brain regions (inner links). The color shade and thickness of the links are proportional to the number of shared GMAS exons between the two tissues.

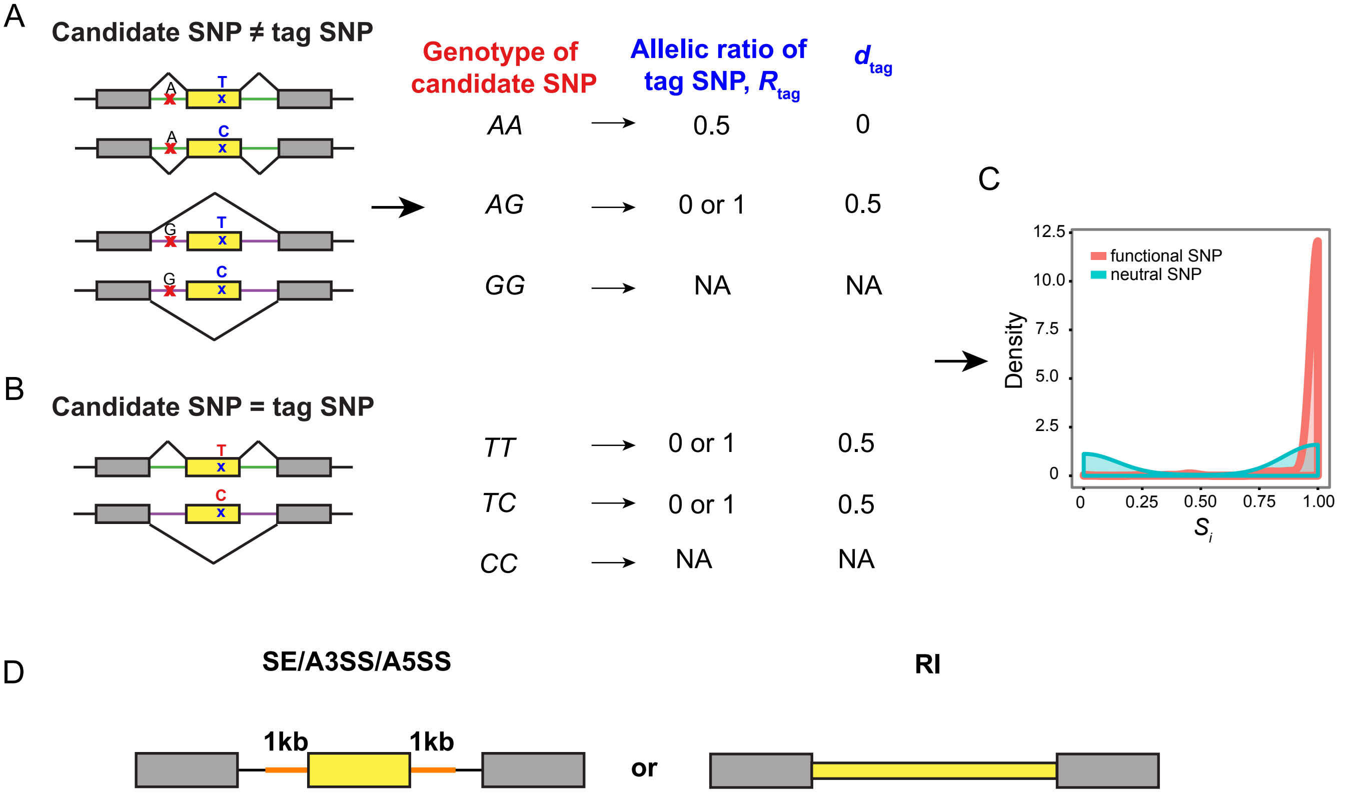


Figure S5. Calculation of concordance scores to predict functional SNPs. (A) A simple example showing a functional candidate SNP that causes complete exon inclusion or exclusion. The allelic ratio of the tag SNP is shown depending on the genotype of the functional SNP. $d_{tag}$ is calculated as $|R_{tag}$-0.5|. (B) A simple example showing that the tag SNP is the functional SNP and its alleles induce complete exon inclusion or exclusion. (C) Hypothetic distribution of concordance score $S_i$ for the functional SNPs in (A) and (B). The distribution for neutral SNPs is also shown. (D) The regions within which to search for candidate functional SNPs (yellow boxes). For SE, A3SS, and A5SS events, we also included intronic SNPs within 1kb (orange line) of the target exon.

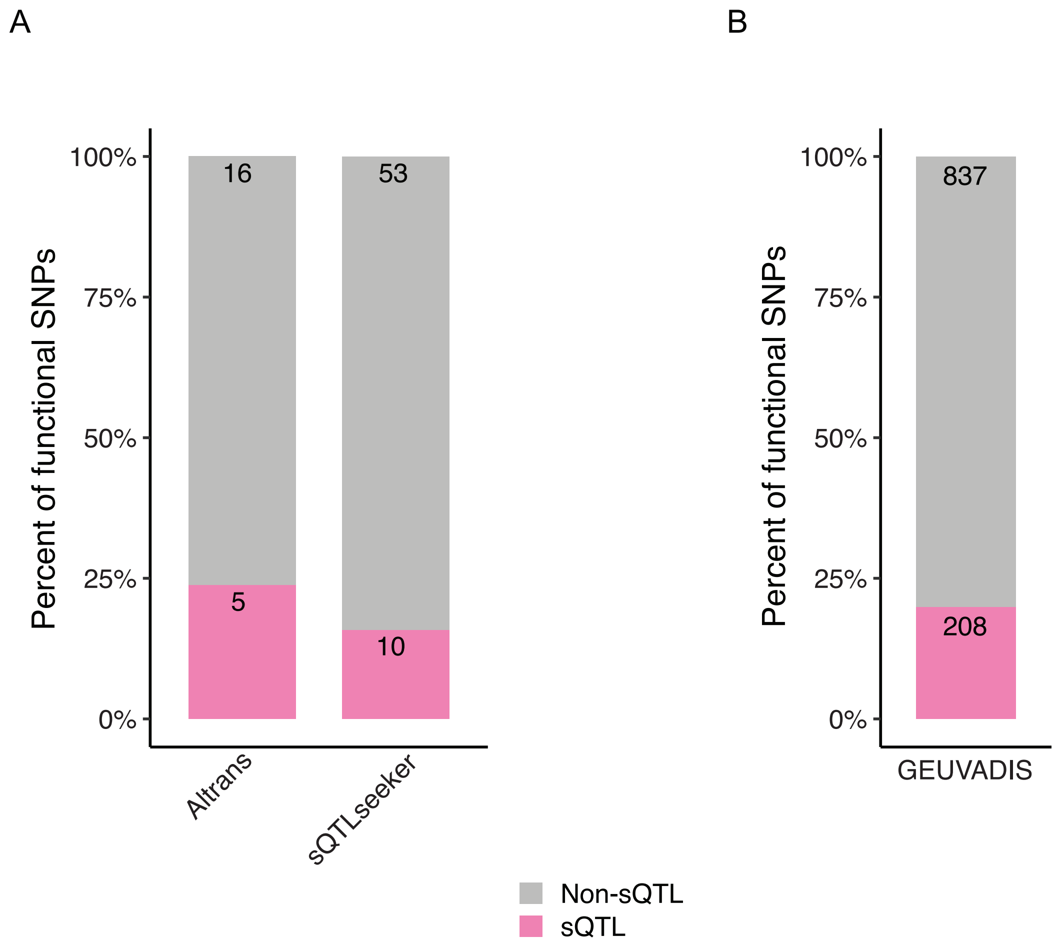


Figure S6. Functional SNPs overlapping known sQTLs. (A) sQTLs identified from GTEx pilot study. (B) sQTLs identified from 1000 Genomes Project.

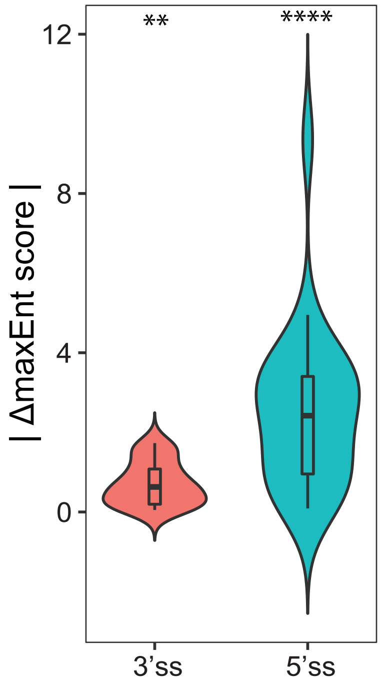


Figure S7. The absolute difference in maxEnt scores between the alternative alleles of predicted functional SNPs located in the 3’ss (Kolmogorov-Smirnov P = 0.003007) or 5‘ss (Kolmogorov-Smirnov P = 0.0001128).

A

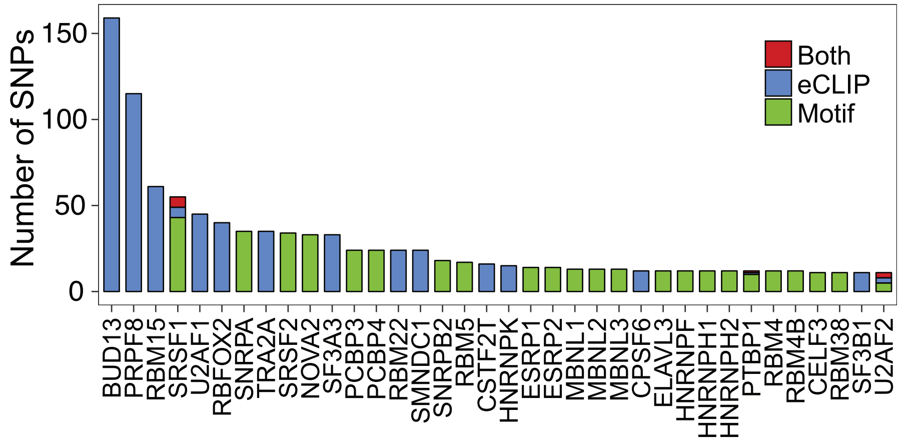


B

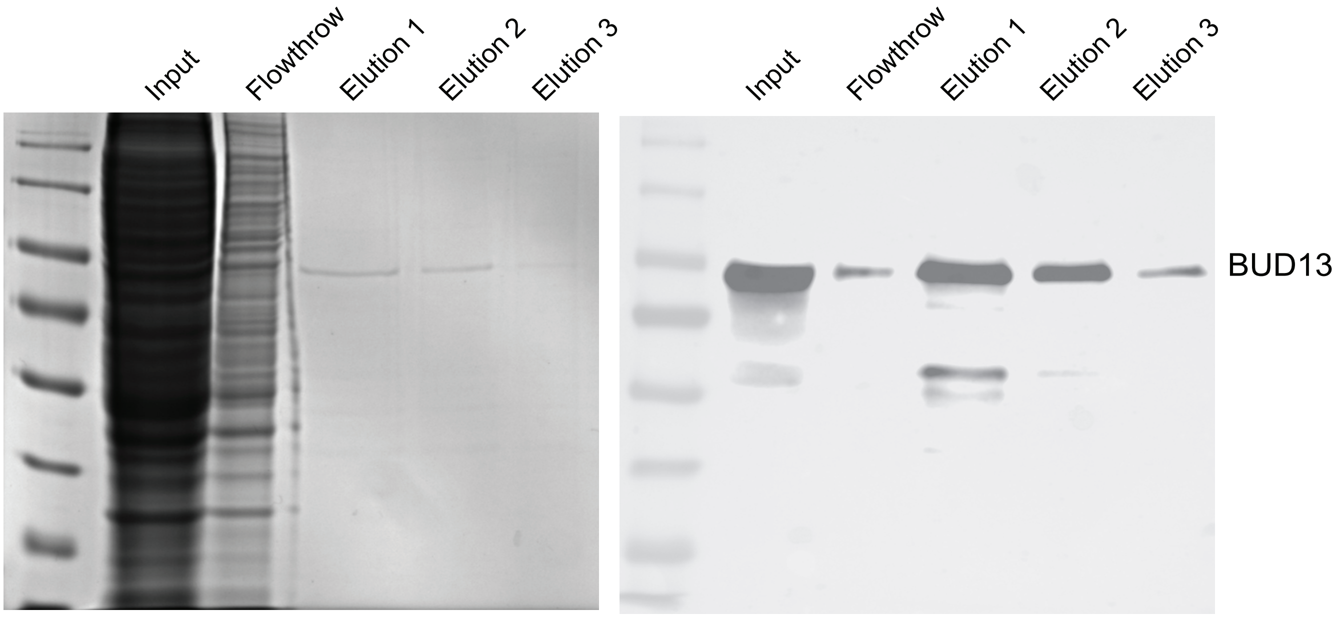


Figure S8. (A) Number of functional SNPs predicted to alter RNA binding motifs of splicing factors by de novo motif and/or eCLIP overlap analyses. Both: functional SNPs predicted by both methods. (B). Purification of recombinant BUD13. Mammalian overexpression of human BUD13 was carried out. pLJM1 3XFlag-Bud13-6HIS lentiviral plasmid was transfected into 293T cells. Stably overexpressed BUD13 was purified using HisTrap purification column and additional Flag affinity purification was performed. Detailed purification conditions are described in Methods. Purified BUD13 proteins were confirmed by SimplyBlue Safe staining (left) and western blot using BUD13 antibody (right).

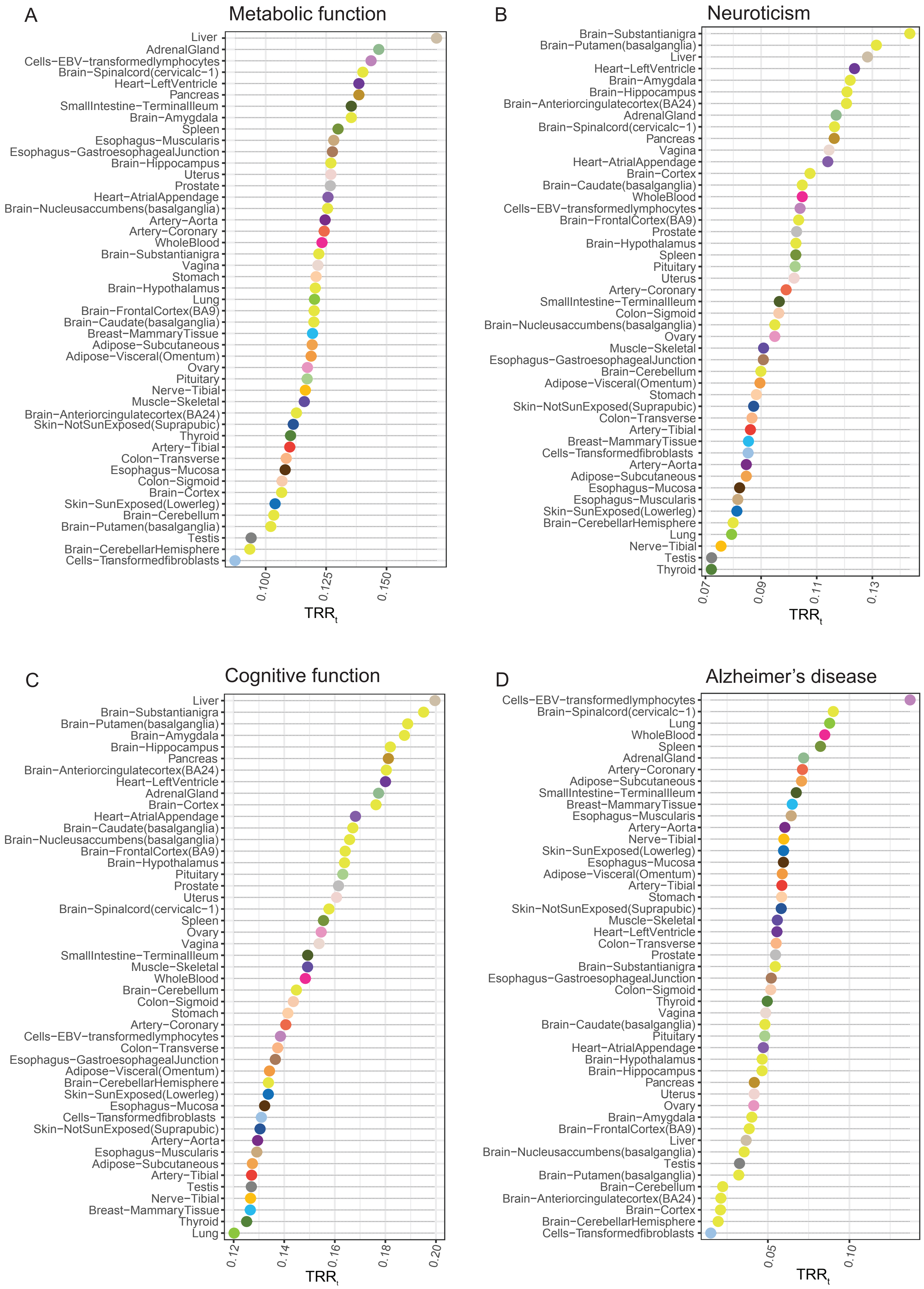

A
Metabolic function
Liver
AdrenalGland
Cells−EBV−transformedlymphocytes
Brain−Spinalcord(cervicalc−1)
Heart−LeftVentricle
Pancreas
SmallIntestine−TerminalIleum
Brain−Amygdala
Spleen
Esophagus−Muscularis
Esophagus−GastroesophagealJunction
Brain−Hippocampus
Uterus
Prostate
Heart−AtrialAppendage
Brain−Nucleusaccumbens(basalganglia)
Artery−Aorta
Artery−Coronary
WholeBlood
Brain−Substantianigra
Vagina
Stomach
Brain−Hypothalamus
Lung
Brain−FrontalCortex(BA9)
Brain−Caudate(basalganglia)
Breast−MammaryTissue
Adipose−Subcutaneous
Adipose−Visceral(Omentum)
Ovary
Pituitary
Nerve−Tibial
Muscle−Skeletal
Brain−Anteriorcingulatecortex(BA24)
Skin−NotSunExposed(Suprapubic)
Thyroid
Artery−Tibial
Colon−Transverse
Esophagus−Mucosa
Colon−Sigmoid
Brain−Cortex
Skin−SunExposed(Lowerleg)
Brain−Cerebellum
Brain−Putamen(basalganglia)
Testis
Brain−CerebellarHemisphere
Cells−Transformedfibroblasts
0.100
0.125
0.150
TRR$_t$
B
Neuroticism
Brain−Substantianigra
Brain−Putamen(basalganglia)
Liver
Heart−LeftVentricle
Brain−Amygdala
Brain−Hippocampus
Brain−Anteriorcingulatecortex(BA24)
AdrenalGland
Brain−Spinalcord(cervicalc−1)
Pancreas
Vagina
Heart−AtrialAppendage
Brain−Cortex
Brain−Caudate(basalganglia)
WholeBlood
Cells−EBV−transformedlymphocytes
Brain−FrontalCortex(BA9)
Prostate
Brain−Hypothalamus
Spleen
Pituitary
Uterus
Artery−Coronary
SmallIntestine−TerminalIleum
Colon−Sigmoid
Brain−Nucleusaccumbens(basalganglia)
Ovary
Muscle−Skeletal
Esophagus−GastroesophagealJunction
Brain−Cerebellum
Adipose−Visceral(Omentum)
Stomach
Skin−NotSunExposed(Suprapubic)
Colon−Transverse
Artery−Tibial
Breast−MammaryTissue
Cells−Transformedfibroblasts
Artery−Aorta
Adipose−Subcutaneous
Esophagus−Mucosa
Esophagus−Muscularis
Skin−SunExposed(Lowerleg)
Brain−CerebellarHemisphere
Lung
Nerve−Tibial
Testis
Thyroid
0.07
0.09
0.11
0.13
TRR$_t$
C
Cognitive function
Liver
Brain−Substantianigra
Brain−Putamen(basalganglia)
Brain−Amygdala
Brain−Hippocampus
Pancreas
Brain−Anteriorcingulatecortex(BA24)
Heart−LeftVentricle
AdrenalGland
Brain−Cortex
Heart−AtrialAppendage
Brain−Caudate(basalganglia)
Brain−Nucleusaccumbens(basalganglia)
Brain−FrontalCortex(BA9)
Brain−Hypothalamus
Pituitary
Prostate
Uterus
Brain−Spinalcord(cervicalc−1)
Spleen
Ovary
Vagina
SmallIntestine−TerminalIleum
Muscle−Skeletal
WholeBlood
Brain−Cerebellum
Colon−Sigmoid
Stomach
Artery−Coronary
Cells−EBV−transformedlymphocytes
Colon−Transverse
Esophagus−GastroesophagealJunction
Adipose−Visceral(Omentum)
Brain−CerebellarHemisphere
Skin−SunExposed(Lowerleg)
Esophagus−Mucosa
Cells−Transformedfibroblasts
Skin−NotSunExposed(Suprapubic)
Artery−Aorta
Esophagus−Muscularis
Adipose−Subcutaneous
Artery−Tibial
Testis
Nerve−Tibial
Breast−MammaryTissue
Thyroid
Lung
0.12
0.14
0.16
0.18
0.20
TRR$_t$
D
Alzheimer's disease
Cells−EBV−transformedlymphocytes
Brain−Spinalcord(cervicalc−1)
Lung
WholeBlood
Spleen
AdrenalGland
Artery−Coronary
Adipose−Subcutaneous
SmallIntestine−TerminalIleum
Breast−MammaryTissue
Esophagus−Muscularis
Artery−Aorta
Nerve−Tibial
Skin−SunExposed(Lowerleg)
Esophagus−Mucosa
Adipose−Visceral(Omentum)
Artery−Tibial
Stomach
Skin−NotSunExposed(Suprapubic)
Muscle−Skeletal
Heart−LeftVentricle
Colon−Transverse
Prostate
Brain−Substantianigra
Esophagus−GastroesophagealJunction
Colon−Sigmoid
Thyroid
Vagina
Brain−Caudate(basalganglia)
Pituitary
Heart−AtrialAppendage
Brain−Hypothalamus
Brain−Hippocampus
Pancreas
Uterus
Ovary
Brain−Amygdala
Brain−FrontalCortex(BA9)
Liver
Brain−Nucleusaccumbens(basalganglia)
Testis
Brain−Putamen(basalganglia)
Brain−Cerebellum
Brain−Anteriorcingulatecortex(BA24)
Brain−Cortex
Brain−CerebellarHemisphere
Cells−Transformedfibroblasts
0.05
0.10
TRR$_t$

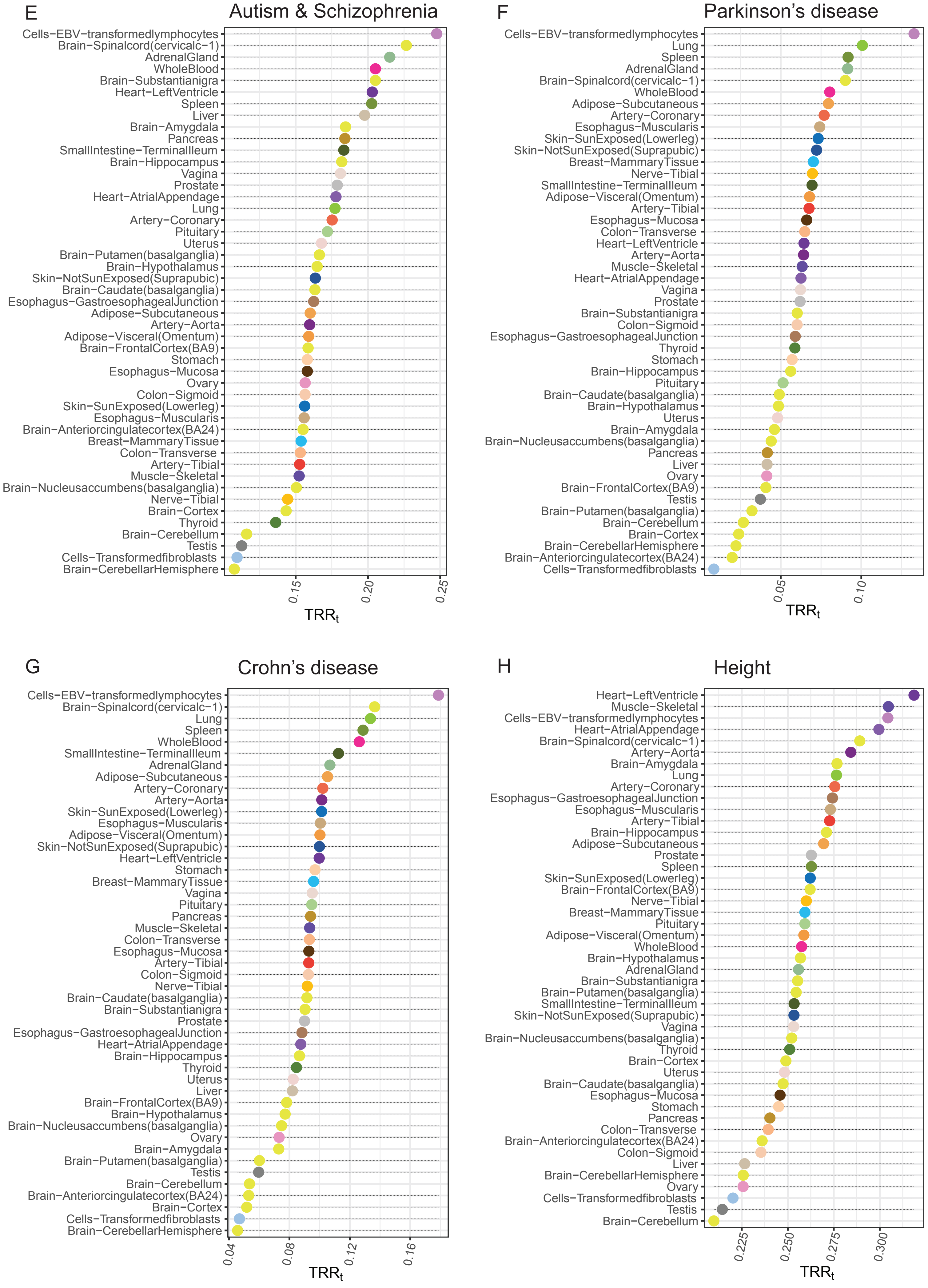
E
Autism & Schizophrenia
Cells−EBV−transformedlymphocytes
Brain−Spinalcord(cervicalc−1)
AdrenalGland
WholeBlood
Brain−Substantianigra
Heart−LeftVentricle
Spleen
Liver
Brain−Amygdala
Pancreas
SmallIntestine−TerminalIleum
Brain−Hippocampus
Vagina
Prostate
Heart−AtrialAppendage
Lung
Artery−Coronary
Pituitary
Uterus
Brain−Putamen(basalganglia)
Brain−Hypothalamus
Skin−NotSunExposed(Suprapubic)
Brain−Caudate(basalganglia)
Esophagus−GastroesophagealJunction
Adipose−Subcutaneous
Artery−Aorta
Adipose−Visceral(Omentum)
Brain−FrontalCortex(BA9)
Stomach
Esophagus−Mucosa
Ovary
Colon−Sigmoid
Skin−SunExposed(Lowerleg)
Esophagus−Muscularis
Brain−Anteriorcingulatecortex(BA24)
Breast−MammaryTissue
Colon−Transverse
Artery−Tibial
Muscle−Skeletal
Brain−Nucleusaccumbens(basalganglia)
Nerve−Tibial
Brain−Cortex
Thyroid
Brain−Cerebellum
Testis
Cells−Transformedfibroblasts
Brain−CerebellarHemisphere
0.15
0.20
0.25
$TRR_t$
F
Parkinson's disease
Cells−EBV−transformedlymphocytes
Lung
Spleen
AdrenalGland
Brain−Spinalcord(cervicalc−1)
WholeBlood
Adipose−Subcutaneous
Artery−Coronary
Esophagus−Muscularis
Skin−SunExposed(Lowerleg)
Skin−NotSunExposed(Suprapubic)
Breast−MammaryTissue
Nerve−Tibial
SmallIntestine−TerminalIleum
Adipose−Visceral(Omentum)
Artery−Tibial
Esophagus−Mucosa
Colon−Transverse
Heart−LeftVentricle
Artery−Aorta
Muscle−Skeletal
Heart−AtrialAppendage
Vagina
Prostate
Brain−Substantianigra
Colon−Sigmoid
Esophagus−GastroesophagealJunction
Thyroid
Stomach
Brain−Hippocampus
Pituitary
Brain−Caudate(basalganglia)
Brain−Hypothalamus
Uterus
Brain−Amygdala
Brain−Nucleusaccumbens(basalganglia)
Pancreas
Liver
Ovary
Brain−FrontalCortex(BA9)
Testis
Brain−Putamen(basalganglia)
Brain−Cerebellum
Brain−Cortex
Brain−CerebellarHemisphere
Brain−Anteriorcingulatecortex(BA24)
Cells−Transformedfibroblasts
0.05
0.10
$TRR_t$
G
Crohn's disease
Cells−EBV−transformedlymphocytes
Brain−Spinalcord(cervicalc−1)
Lung
Spleen
WholeBlood
SmallIntestine−TerminalIleum
AdrenalGland
Adipose−Subcutaneous
Artery−Coronary
Artery−Aorta
Skin−SunExposed(Lowerleg)
Esophagus−Muscularis
Adipose−Visceral(Omentum)
Skin−NotSunExposed(Suprapubic)
Heart−LeftVentricle
Stomach
Breast−MammaryTissue
Vagina
Pituitary
Pancreas
Muscle−Skeletal
Colon−Transverse
Esophagus−Mucosa
Artery−Tibial
Colon−Sigmoid
Nerve−Tibial
Brain−Caudate(basalganglia)
Brain−Substantianigra
Prostate
Esophagus−GastroesophagealJunction
Heart−AtrialAppendage
Brain−Hippocampus
Thyroid
Uterus
Liver
Brain−FrontalCortex(BA9)
Brain−Hypothalamus
Brain−Nucleusaccumbens(basalganglia)
Ovary
Brain−Amygdala
Brain−Putamen(basalganglia)
Testis
Brain−Cerebellum
Brain−Anteriorcingulatecortex(BA24)
Brain−Cortex
Cells−Transformedfibroblasts
Brain−CerebellarHemisphere
0.04
0.08
0.12
0.16
$TRR_t$
H
Height
Heart−LeftVentricle
Muscle−Skeletal
Cells−EBV−transformedlymphocytes
Heart−AtrialAppendage
Brain−Spinalcord(cervicalc−1)
Artery−Aorta
Brain−Amygdala
Lung
Artery−Coronary
Esophagus−GastroesophagealJunction
Esophagus−Muscularis
Artery−Tibial
Brain−Hippocampus
Adipose−Subcutaneous
Prostate
Spleen
Skin−SunExposed(Lowerleg)
Brain−FrontalCortex(BA9)
Nerve−Tibial
Breast−MammaryTissue
Pituitary
Adipose−Visceral(Omentum)
WholeBlood
Brain−Hypothalamus
AdrenalGland
Brain−Substantianigra
Brain−Putamen(basalganglia)
SmallIntestine−TerminalIleum
Skin−NotSunExposed(Suprapubic)
Vagina
Brain−Nucleusaccumbens(basalganglia)
Thyroid
Brain−Cortex
Uterus
Brain−Caudate(basalganglia)
Esophagus−Mucosa
Stomach
Pancreas
Colon−Transverse
Brain−Anteriorcingulatecortex(BA24)
Colon−Sigmoid
Liver
Brain−CerebellarHemisphere
Ovary
Cells−Transformedfibroblasts
Testis
Brain−Cerebellum
0.225
0.250
0.275
0.300
$TRR_t$

I

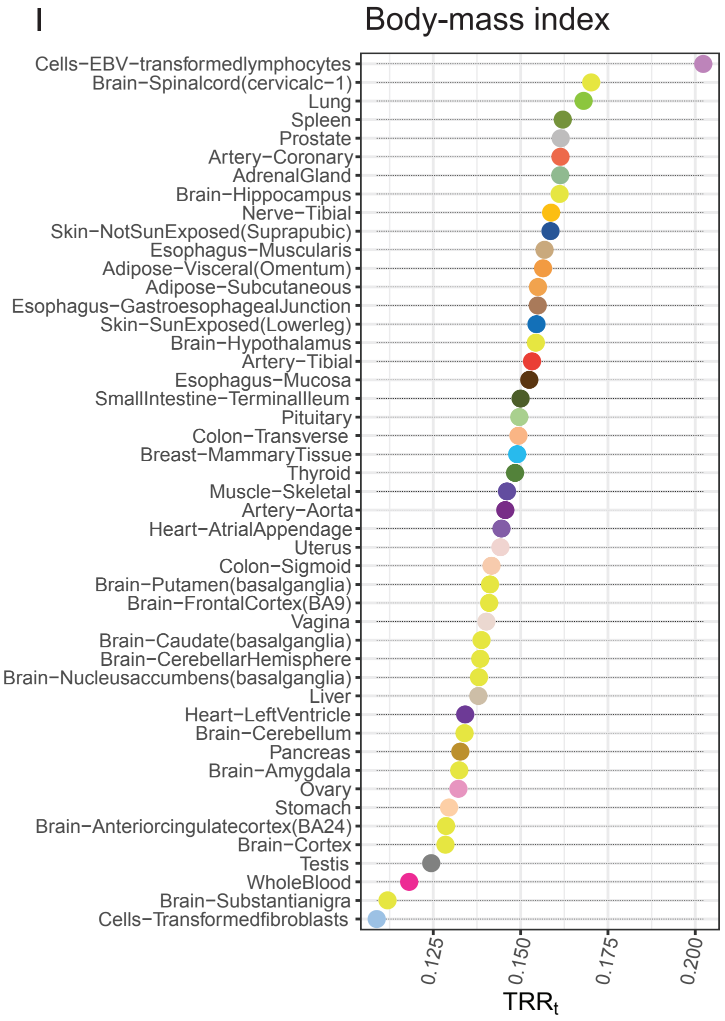


Figure S9. (A)-(I)Trait relevance ratios for all GMAS SNPS in LD with GWAS SNPs for different traits/diseases, respectively.
